# Radial F-actin Organization During Early Neuronal Development

**DOI:** 10.1101/372813

**Authors:** Durga Praveen Meka, Robin Scharrenberg, Bing Zhao, Theresa König, Irina Schaefer, Birgit Schwanke, Oliver Kobler, Sergei Klykov, Melanie Richter, Dennis Eggert, Sabine Windhorst, Carlos G. Dotti, Michael R. Kreutz, Marina Mikhaylova, Froylan Calderon de Anda

**Affiliations:** RG Neuronal Development, Center for Molecular Neurobiology Hamburg (ZMNH), University Medical Center Hamburg-Eppendorf, 20251 Hamburg, Germany.; Combinatorial Neuroimaging Core Facility (CNI), Leibniz Institute for Neurobiology, 39118 Magdeburg, Germany.; Emmy-Noether Group “Neuronal Protein Transport”, Center for Molecular Neurobiology (ZMNH), University Medical Center Hamburg-Eppendorf, 20251 Hamburg, Germany.; Max Planck Institute for the Structure and Dynamics of Matter, 22761 Hamburg and Heinrich Pette Institute - Leibniz Institute for Experimental Virology, 20251 Hamburg, Germany.; Department of Biochemistry and Signal Transduction, University Medical Center Hamburg-Eppendorf, 20246 Hamburg, Germany.; Centro de Biología Molecular “Severo Ochoa”, CSIC-UAM, Madrid, Spain.; RG Neuroplasticity, Leibniz Institute for Neurobiology, 39118 Magdeburg, Germany.; Leibniz Guest Group “Dendritic Organelles and Synaptic Function”, Center for Molecular Neurobiology (ZMNH), University Medical Center Hamburg-Eppendorf, 20251 Hamburg, Germany.

## Abstract

The centrosome is thought to be the major neuronal microtubule-organizing center (MTOC) in early neuronal development, producing microtubules with a radial organization. In addition, albeit in vitro, recent work showed that isolated centrosomes could serve as an actin-organizing center (Farina et al., 2016), raising the possibility that neuronal development may, in addition, require a centrosome-based actin radial organization. Here we report, using super-resolution microscopy and live-cell imaging, F-actin organization around the centrosome with dynamic F-actin aster-like structures with F-actin fibers extending and retracting actively. Photoconversion/photoactivation experiments and molecular manipulations of F-actin stability reveal a robust flux of somatic F-actin towards the cell periphery. Finally, we show that somatic F-actin intermingles with centrosomal PCM-1 satellites. Knockdown of PCM-1 and disruption of centrosomal activity not only affect F-actin dynamics near the centrosome but also in distal growth cones. Collectively the data show a radial F-actin organization during early neuronal development, which might be a cellular mechanism for providing peripheral regions with a fast and continuous source of actin polymers; hence sustaining initial neuronal development.

## Introduction

The centrosome is thought to be the major neuronal microtubule-organizing center (MTOC) in early developing neurons (Calderon de Anda et al., 2008; Stiess et al., 2010; Yau et al., 2014), producing microtubules with a radial organization (Sakakibara et al., 2014; Yau et al., 2016). Recently, it has been shown that isolated centrosomes can serve as an actin-organizing center *in vitro* (Farina et al., 2016), suggesting that the centrosome might control F-actin organization and dynamics during initial neuronal development. However, initial attempts to demonstrate that somatic F-actin can be delivered rapidly to distal growth cones were not successful (Bernstein and Bamburg, 1992; Sanders and Wang, 1991). Moreover, the classical view on the role of actin on neuronal development is contrary to this idea. For instance, numerous studies have demonstrated that F-actin is assembled locally in growth cones and that impaired local assembly is sufficient to block neurite growth (Flynn et al., 2012; Forscher et al., 1992; Gallo et al., 2002; Lowery and Van Vactor, 2009; Okabe and Hirokawa, 1991). Nevertheless, several other studies have reported that growth cone-like structures, comprised of F-actin, have an anterograde wave-like propagation along neurites, supporting neurite extension (Flynn et al., 2009; Ruthel and Banker, 1998, 1999; Winans et al., 2016). Other studies described anterograde F-actin flow during neuronal migration (He et al., 2010; Solecki et al., 2009) and at the base of growth cones (Burnette et al., 2008). Thus, adding weight to the possibility that centrifugal actin forces starting in the cell body may contribute to the final neuronal phenotype during development. To test this possibility, we performed a series of state-of-the-art methods to examine somatic F-actin organization and dynamics in living neurons.

Additionally, we propose PCM-1 as a molecular determinant for somatic F-actin organization. PCM-1 has been shown to promote F-actin polymerization in non neuronal cells (Farina et al., 2016) and PCM-1-containing pericentriolar satellites are important for the recruitment of proteins that regulate centrosome function (Dammermann and Merdes, 2002). The depletion of PCM-1 disrupts the radial organization of microtubules without affecting microtubule nucleation (Dammermann and Merdes, 2002). PCM-1 particles preferentially localize near the centrosome (de Anda et al., 2010; Ge et al., 2010). In previous work we found PCM-1 down-regulation in the developing cortex to disrupt neuronal polarization and to preclude axon formation (de Anda et al., 2010). Furthermore, neuronal migration was impaired with piling up of neurons in the intermediate zone (de Anda et al., 2010). Here, we demonstrate that PCM-1 determines not only the somatic F-actin organization and dynamics but also that lack of PCM-1 has a radial effect, which ultimately disrupts growth cone dynamics and neurite length. Overall, our data show a novel somatic F-actin organization, which regulates early neuronal development *in vitro* and *in vivo*.

### Results F-actin organizes around the centrosome

We studied the micro and nano-structural organization of cytosolic F-actin near the centrosome via confocal and super-resolution microscopy during early neuronal differentiation *in vitro* (from stage 1 to early stage 3; (Dotti et al., 1988)) and *in situ*. To this end the F-actin cytoskeleton in fixed and live cells was visualized via confocal and STED microscopy by labeling cells with Phalloidin-488, Phalloidin Atto647N, SiR-actin probe or Lifeact-GFP (Lukinavicius et al., 2014). Confocal microscopy showed a preferential localization of cytosolic F-actin puncta near the centrosome in cultured neurons labeled with phalloidin and in neurons in the developing cortex labeled with Lifeact-GFP (Fig. 1A-C and Fig. S1A-C). STED microscopy images revealed that somatic F-actin organized as tightly packed structures constituted by a core of dense F-actin attached to F-actin fibers (aster-like structures, Fig. 1D and Fig. S1D). Moreover, we used single molecule localization microscopy (SMLM / STORM) of Phalloidin-Alexa647 labeled F-actin and corroborated that it organized around the centrosome in a pocket-like structure, where several F-actin puncta surrounded the centrosome with individual puncta exhibiting an aster-like organization (Fig. S2).

**Fig. 1.**
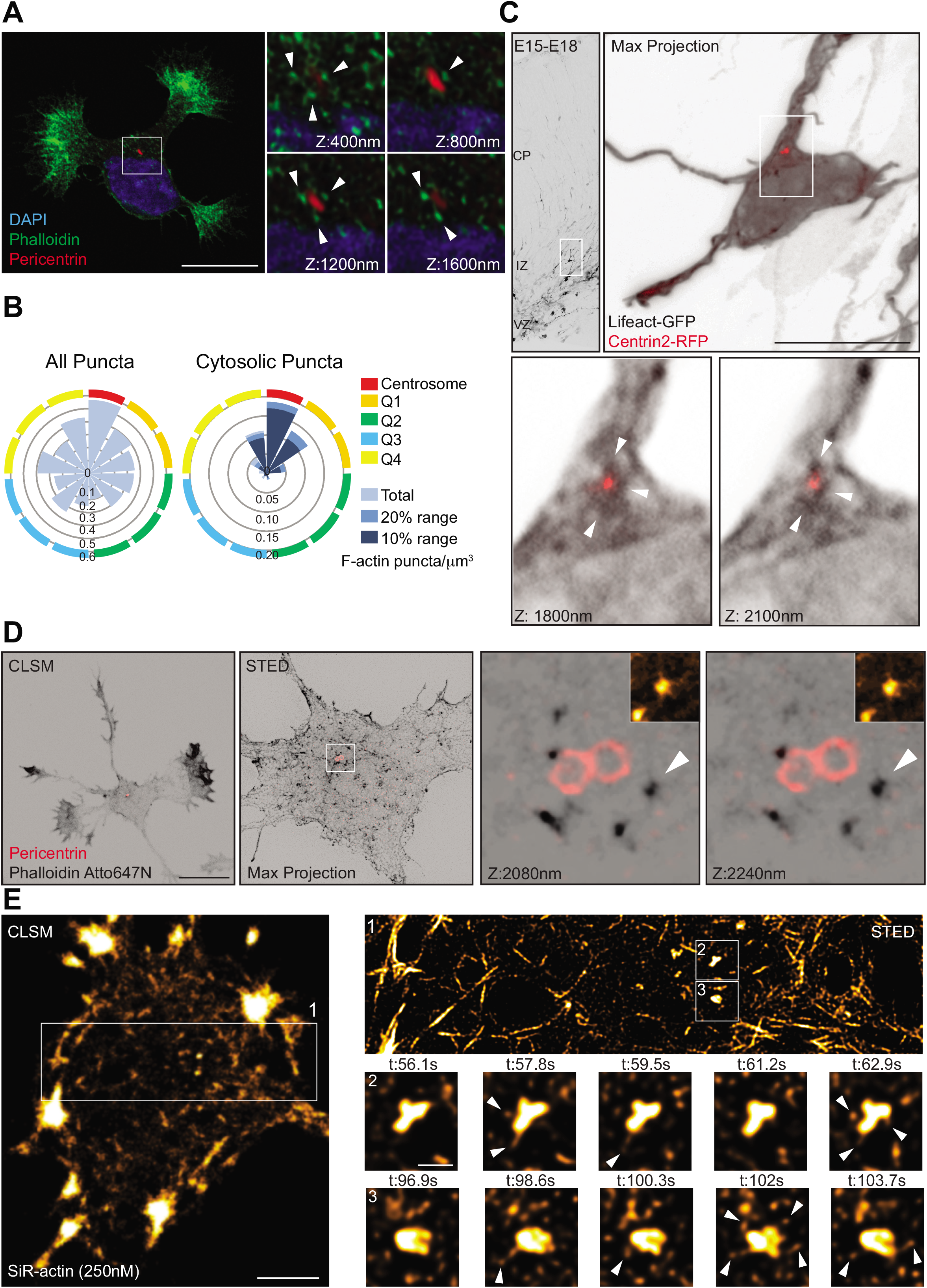
Super-resolution microscopy reveals cytosolic F-actin puncta releasing F-actin fibers in developing neurons. **(A)** Stage 2 hippocampal neuron labelled with phalloidin and Pericentrin antibody, confocal z-stacks from the inset show F-actin puncta around the centrosome. **(B)** Rose plots depict distribution of F-actin puncta (cortical and cytosolic or only cytosolic) in the cell body with respect to the position of the centrosome. 10% and 20% distance range from the total area is indicated in different shades of blue; n=12 cells from at least three different cultures. **(C)** Multipolar cell located in the IZ of the developing cortex expresses Lifeact-GFP and Centrin2-RFP and shows F-actin puncta surrounding the centrosome in the developing cortex. **(D)** Confocal (CLSM) and STED microscopic images of stage 2 hippocampal neuron. Inset: STED Z-stack images with 160 nm Z-spacing showing F-actin puncta localizing near the centrosome. Insets from arrowheads in Z-stack images show F-actin puncta with F-actin fibers attached. **(E)** Confocal image of stage 2 neuron labeled with SiR-actin (250nM). Inset 1: snap shot of STED time-lapse. Time-lapse montages from insets 2 and 3 depict individual F-actin puncta releasing F-actin fibers (arrowheads). Time (t) interval = 1.7 sec (Movie S1). Scale bar: 10 μm (A, C and D); 2 μm (E); 0.5 μm (inset E).

In order to determine whether or not somatic F-actin puncta represent true sites of actin polymerization, we transfected cells with Lifeact-GFP and performed epifluorescence time-lapse imaging (frame rate: 2 sec for 5 min) on DIV1 neurons. Cells expressing high levels of Lifeact showed stabilized F-actin in the form of somatic F-actin fibers (Fig. S3). Therefore, we exclusively analyzed cells where Lifeact expression labeled the F-actin cytoskeleton at similar levels as detected with phalloidin staining (Fig. S3). Time-lapse analysis of cells co-transfected with Lifeact-GFP and the microtubule plus-end marker EB3-mCherry corroborated that F-actin puncta concentrate near the MTOC identified by radial trajectories of EB3-mCherry comets (Fig. S4A, B). Moreover, our recordings showed that the F-actin puncta in the soma are highly dynamic and intermittent. These puncta exhibit a repetitive appearance and disappearance at the same location as shown via kymographs (Fig. S4C, D). Based on the duration of appearance, we categorized them as unstable (<15 sec), intermediately stable (16-240 sec), and long-lasting (241-300 sec) F-actin puncta. The majority of puncta are unstable (Fig. S4E), suggesting that these puncta are places of high F-actin turnover. Accordingly, our FRAP analysis of somatic F-actin puncta showed fast fluorescence recovery (Fig. S5A, B); thus, confirming that somatic F-actin puncta are places of high F-actin turnover. We also found that these F-actin puncta in the cell body release F-actin comets (pointed by red arrowheads in Fig. S4B), which suggest that they might function as a source of somatic F-actin.

To gain further insight into the relevance of this F-actin organization, we employed STED time-lapse microscopy labeling F-actin with SiR-actin. Since SiR-actin is known to stabilize F-actin, we tested different concentrations of SiR-actin (250 nM and 500 nM). Cells labeled with 500 nM SiR-actin showed less dynamic F-actin aster-like structures, with longer F-actin fibers attached to the F-actin core (Fig. S6). Whereas, cells labeled with 250 nM SiR-actin showed aster-like F-actin structures that are highly dynamic, extending and retracting F-actin fibers constantly in the range of seconds (Fig. 1E; Movie S1). Altogether, these results unveil the existence of a complex and dynamic somatic F-actin organization near the centrosome, suggesting a role in neuronal development.

### Somatic F-actin is radially delivered to the cell periphery

We therefore asked whether somatic actin polymerization could serve as a source for cell peripheral F-actin. To this end, we used DIV 1 neurons transfected with Lifeact-mEos3.2, which undergoes an irreversible photoconversion in response to 405 nm light from green to red fluorescence with emission peaks at 516 nm to 581 nm respectively. We had to irradiate several F-actin puncta (5.2 to 7.1 μm^2^) at once given that single punctum irradiation (2.2 μm^2^) did not yield enough traceable converted signal when spreading further (Fig. S7A). Interestingly, when we photoconverted a group of F-actin puncta in the soma, the intensity of the converted F-actin puncta decreased with time concomitant with a fast increase of converted signal in the cell periphery/growth cones (Fig. 2A, B and Movie S2). Another actin probe (actin-mEos4b), which labels F-actin and actin monomers, also distributed into growth cones after irradiation (Fig. 2B; Fig. S7B; Movie S4). However, irradiated mEos3.2 alone resulted in reduced movement of the probe as well as decreased enrichment in growth cones compared to Lifeact-mEos3.2 (Fig. 2B; Fig. S7C; Movie S4).

**Fig. 2.**
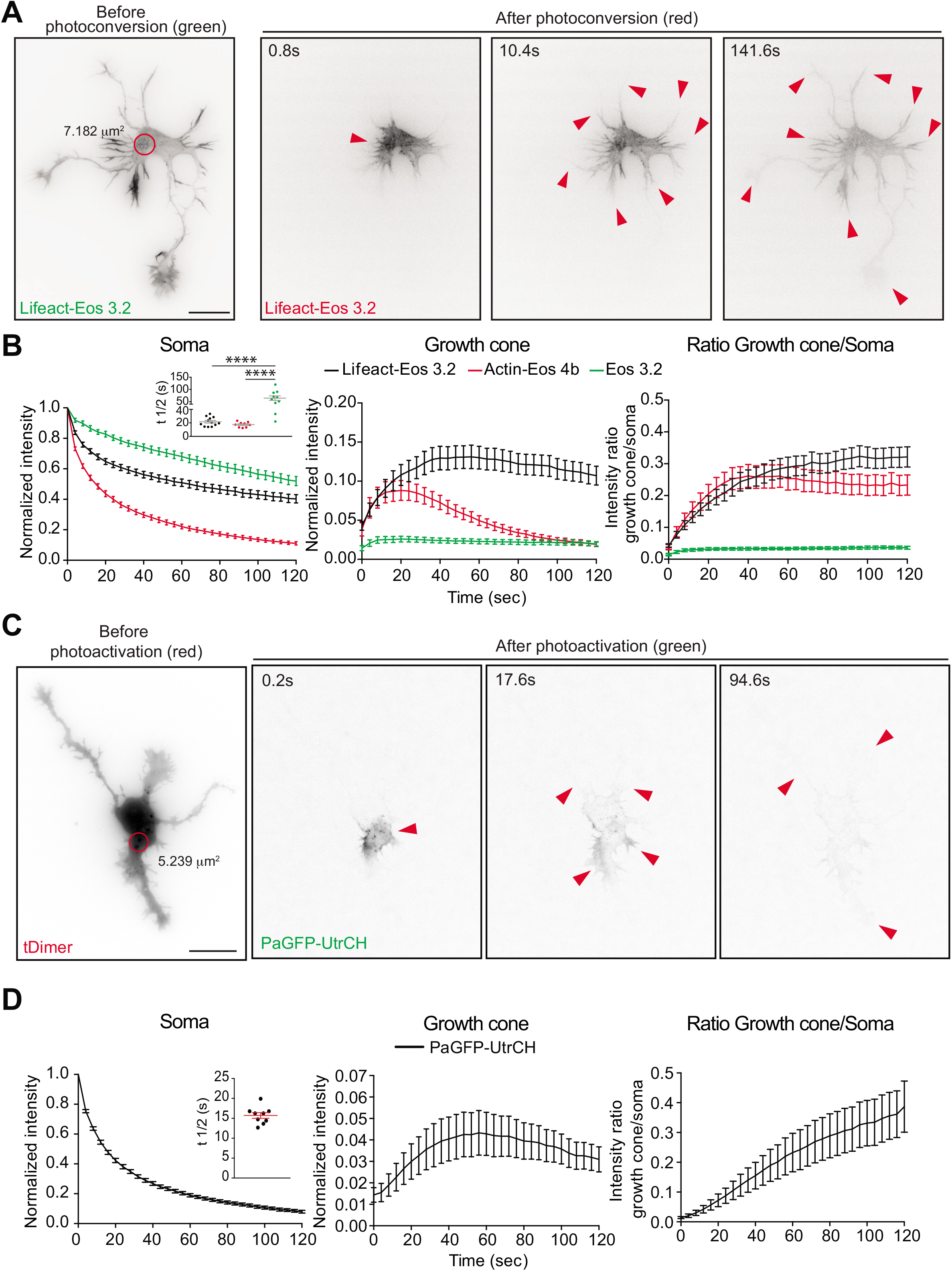
Somatic F-actin puncta act as rapid supply sources of F-actin to the periphery in developing neurons. **(A)** Lifeact-mEos3.2-expressing stage 2 neuron photoconverted in the soma with 405 nm laser (red circle). Cell before (green signal) and after photoconversion (red signal); red arrowheads point reach of the photoconverted signal over time. **(B)** Left panel: normalized intensities in the photoconverted area of Lifeact-mEos3.2, Actin-mEos4b or mEos3.2 expressing cells. Inset graph: Half-time (t ½) values for Lifeact-mEos3.2 = 21.66 ± 1.937, Actin-mEos4b = 17.40 ± 1.275 and mEos3.2 = 64.65 ± 9.205. p<0.0001 by one-way ANOVA, post hoc Dunnett’s test, ***p<0.001, Mean ± SEM. Middle panel: photoconverted signal in growth cones over time relative to the average initial signal from irradiated area for Lifeact-mEos3.2, Actin-mEos4b or mEos3.2 expressing cells. Right panel: Ratio of signal in Growth cone compared to soma upon photoconversion over time; All panels: n=12 cells for Lifeact-mEos3.2, n=11 cells for mEos3.2 and n=9 cells for Actin-mEos4b. **(C)** Neuron before (red; tDimer) and after photoactivation (green; PaGFP-UtrCH) in the soma with 405 nm laser (red circle). Red arrowheads point the reach of the photoactivated signal over time. **(D)** Left panel: normalized intensity values in the photoactivated area of PaGFP-UtrCH expressing cells. Inset: half-time (t ½) values for PaGFP-UtrCH = 15.71 ± 0.712, Mean ± SEM, n=9 cells. Middle panel: photoconverted signal in growth cones over time relative to the average initial signal from irradiated area for PaGFP-UtrCH expressing cells. Right panel: Growth cone to soma photoactivated signal intensity ratio of PaGFP-UtrCH expressing cells. Scale bar: 10 μm (A, C).

To confirm that the radial translocation of Lifeact signal is due to the movement of F-actin but not actin monomers bound to Lifeact, we treated the cells with Cytochalasin D, which disrupts the F-actin cytoskeleton. It has been shown that Cytochalasin D treatment induces F-actin clusters around the centrosome in nonneuronal cells (Farina et al., 2016). Similarly, we observed that Cytochalasin D treatment in neurons also induced the formation of F-actin aggregates near the centrosome from pre-existing intermittent F-actin puncta (Fig. S8A, B). These clusters do not depend on membrane organization since brefeldin A, which disrupts Golgi and endoplasmic reticulum, did not affect the localization of F-actin clusters near the centrosome (Fig. S8C). Photoconversion of the somatic F-actin clusters of cytochalasin D treated cells did not induce Lifeact translocation towards the cell periphery (Fig. S8D, E and Movie S5), indicating that the translocation of photoconverted signal is not due to the movement of the Lifeact bound to actin monomers but indeed labeled F-actin. Although Lifeact binds *in vitro* with higher affinity to actin monomers than to F-actin (Riedl et al., 2008), it is still possible that in cells the amount of Lifeact bound to actin monomers is not the predominant species. This is also suggested by the pattern of Lifeact expression, which resembles the Phalloidin staining (Fig. S3).

Notably, we also found that the somatic photoactivation of Photoactivatable GFP-Utrophin (PaGFP-UtrCH), which specifically labels F-actin (Burkel et al., 2007; Melak et al., 2017), leads to the distribution of photoactivated PaGFP-UtrCH in the cell periphery (Fig. 2C, D and Movie S3) similarly as Lifeact-GFP. Therefore, our results demonstrate that filamentous actin translocates from the soma to the cell periphery.

Further characterization of photoconverted Lifeact-mEos3.2 (red signal) or photoactivatable PaGFP-UtrCH in the cell periphery showed that translocation does not occur preferentially to the growth cone of the longest neurite (Fig. 3A, B) but to the growth cone containing more F-actin (green arrow in Fig. 3C, D).

**Fig. 3.**
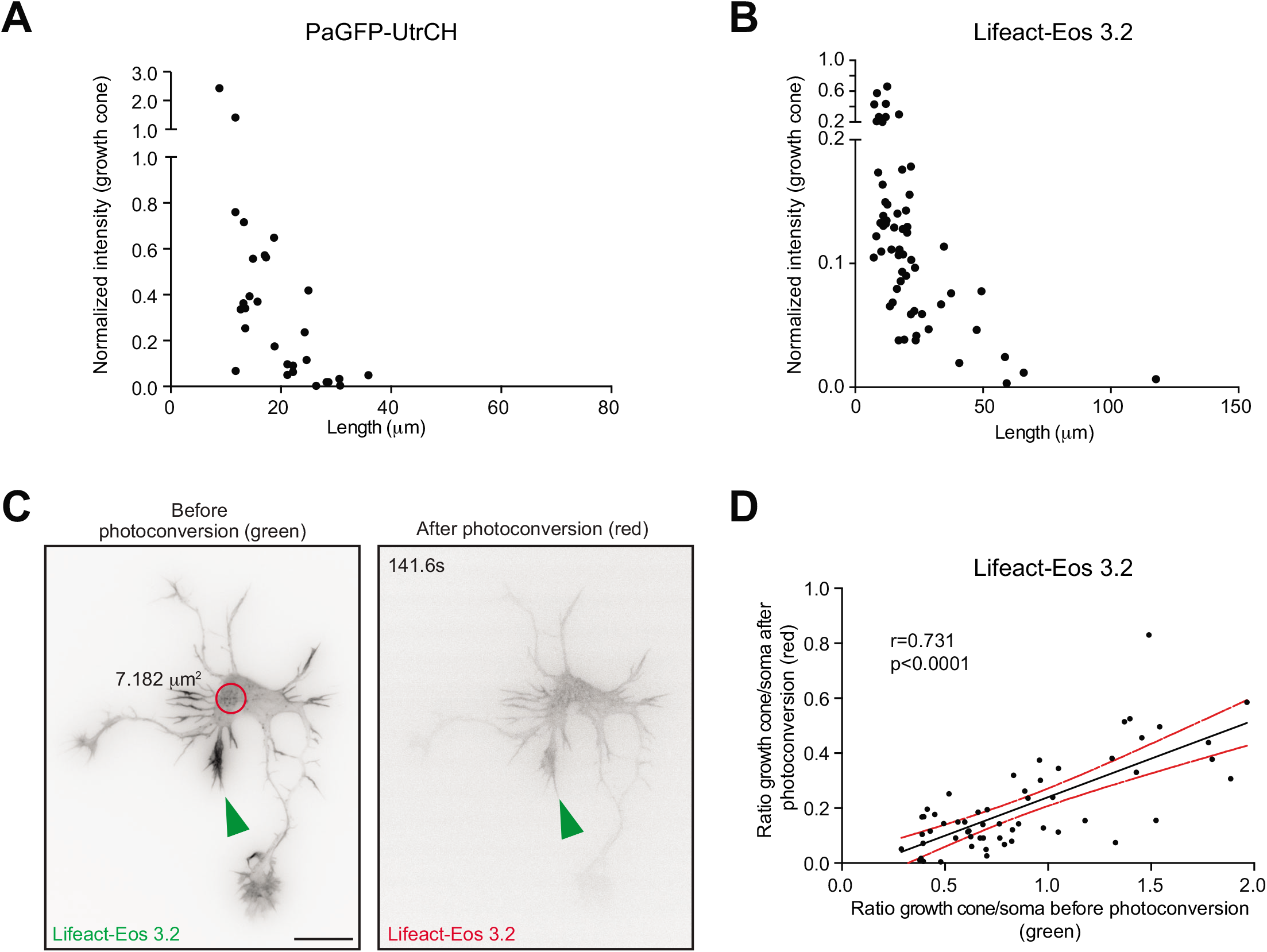
Somatic F-actin translocation occurs preferentially to the growth cone with higher F-actin content. **(A, B)** Normalized intensity values from the growth cones plotted against their neurite lengths of **(A)** PaGFP-UtrCH or **(B)** Lifeact-mEos3.2 expressing cells 120 sec after photoactivation (PaGFP-UtrCH) or photoconversion (Lifeact-mEos3.2). **(C)** Cell from figure 2 showing the preferential translocation of photoconverted Lifeact-mEos3.2 to the growth cone with higher content of Lifeact-mEos3.2 before photoconversion (green arrowhead). **(D)** Pearson correlation of growth cone to soma intensity ratio before and after photoconversion in Lifeact-mEos3.2 expressing cells (n=57 neurites; Pearson r = 0.731, p<0.0001). Scale bar: 10 μm (A).

Next, we tested whether or not F-actin translocation is exclusively radially oriented. Consequently, we decided to irradiate growth cones labeled with Lifeact-mEos3.2, PaGFP-UtrCH, Actin-mEos4b, or mEos3.2. When mEos3.2 or Actin-mEos4b transfected neurons were irradiated at growth cones, the converted signal translocated towards the cell body (Fig. 4B; Fig. S9B, C; Movie S6). In contrast, irradiation of growth cones labeled with Lifeact-mEos3.2 or PaGFP-UtrCH did not induce retrograde movement of photoconverted Lifeact-mEos3.2 or photoactivated PaGFP-UtrCH signal to the cell body (Fig. 4, Fig. S9A; Movie S6); thus, showing that F-actin translocation is unidirectional.

**Fig. 4.**
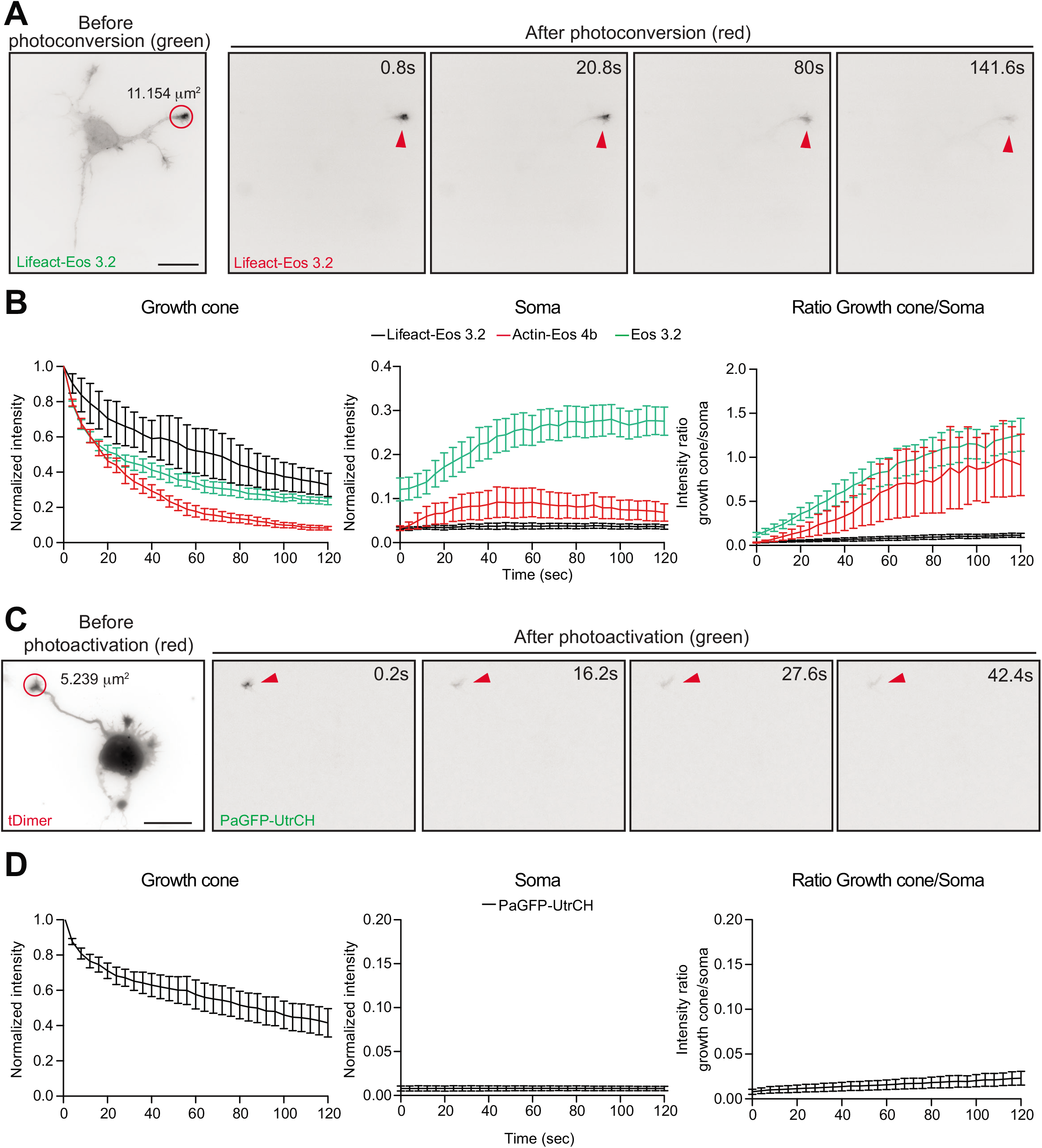
F-actin translocation is exclusively radially oriented. **(A)** Lifeact-mEos3.2 expressing cell before (green) and after photoconversion (red) at the growth cone with 405 nm laser (red circle). Red arrowheads point the reach of the photoconverted signal over time. **(B)** Left panel: normalized intensity values in the photoconverted area (growth cones) of Lifeact-mEos3.2, Actin-mEos4b or mEos3.2 expressing cells. Middle panel: photoconverted signal in the soma over time relative to the average initial signal from irradiated area for Lifeact-mEos3.2, Actin-mEos4b and mEos3.2 expressing cells. Right panel: growth cone to soma photoconverted signal intensity ratio of Lifeact-mEos3.2, Actin-mEos4b and mEos3.2 expressing cells. Mean ± SEM, n=3 cells for Lifeact-mEos3.2, n=3 cells for Actin-mEos4b, and n=7 cells for mEos3.2 from two different cultures. **(C)** Neuron before (red; tDimer) and after photoactivation (green; PaGFP-UtrCH) at the growth cone with 405 nm laser (red circle). Red arrowheads point the reach of the photoactivated signal over time. **(D)** Left panel: normalized intensity values in the photoactivated area (growth cones) of PaGFP-UtrCH expressing cells. Middle panel: photoconverted signal in the soma over time relative to the average initial signal from irradiated area for PaGFP-UtrCH expressing cells. Right panel: Growth cone to soma intensity ratio of photoactivated UtrCH signal. Mean ± SEM, n=8. Scale bar: 10 μm (A, D).

In order to decrease F-actin dynamics to better resolve the somatic F-actin translocation to the cell periphery, we treated our cultured neurons with Jasplakinolide, an agent which stabilizes polymerized actin filaments and stimulates actin filament nucleation, (Bubb et al., 2000). However, our treatment (0.3, 0.5 and 1 microM / hr) precluded the existence of peripheral F-actin and induced the formation of a somatic F-actin ring-structure, detected both with Lifeact and Phalloidin staining (Fig. S10A, B). This structure appeared in the area with the highest density of plus-end microtubules labeled via EB3-mCherry (Fig. S10B). These results suggest that Jasplakinolide prevents F-actin dynamics, thus, “locking” the F-actin in the soma and blocking its movement. Photoconversion experiments further confirm these findings showing no radial movement of photoconverted signal to the periphery in Jasplakinolide-treated cells (Fig. S10C, D).

Importantly, with Drebrin or Cofilin constructs, as F-actin stabilizing tools, we could decrease the overall dynamics of F-actin without completely stabilizing/freezing the actin filaments. Drebrin inhibits Cofilin-induced severing of F-actin and stabilizes F-actin (Grintsevich and Reisler, 2014; Mikati et al., 2013); Drebrin phosphorylation at S142 promotes F-actin bundling (Worth et al., 2013). Therefore, the Drebrin phosphomimetic mutant (S142D) is a suitable candidate to decrease overall F-actin dynamics. Similarly, phosphomimetic Cofilin (S3E) is not able to sever F-actin, thus it decreases F-actin turnover (Chai et al., 2009). Previously, it was shown that Drebrin co-localizes with F-actin in growth cones (Geraldo et al., 2008). Time-lapse microscopy analysis of Drebrin transfected cells revealed Drebrin to co-localize with F-actin puncta in the cell body (Fig. 5A; Movie S7). Moreover, transfection with the phospho-mimetic mutant Drebrin-S142D reduced F-actin treadmilling compared to cells expressing only Lifeact-GFP (Fig. S11A and (Zhao et al., 2017)). Importantly, the total number of somatic F-actin puncta decreased after Drebrin-S142D expression (Fig. 5D) with an increase in the relative number of long-lasting F-actin puncta (Fig. 5E). Interestingly, the stable F-actin puncta released noticeable F-actin comet-like structures towards the cell periphery (Fig. 5B, Movie S7).

**Fig. 5.**
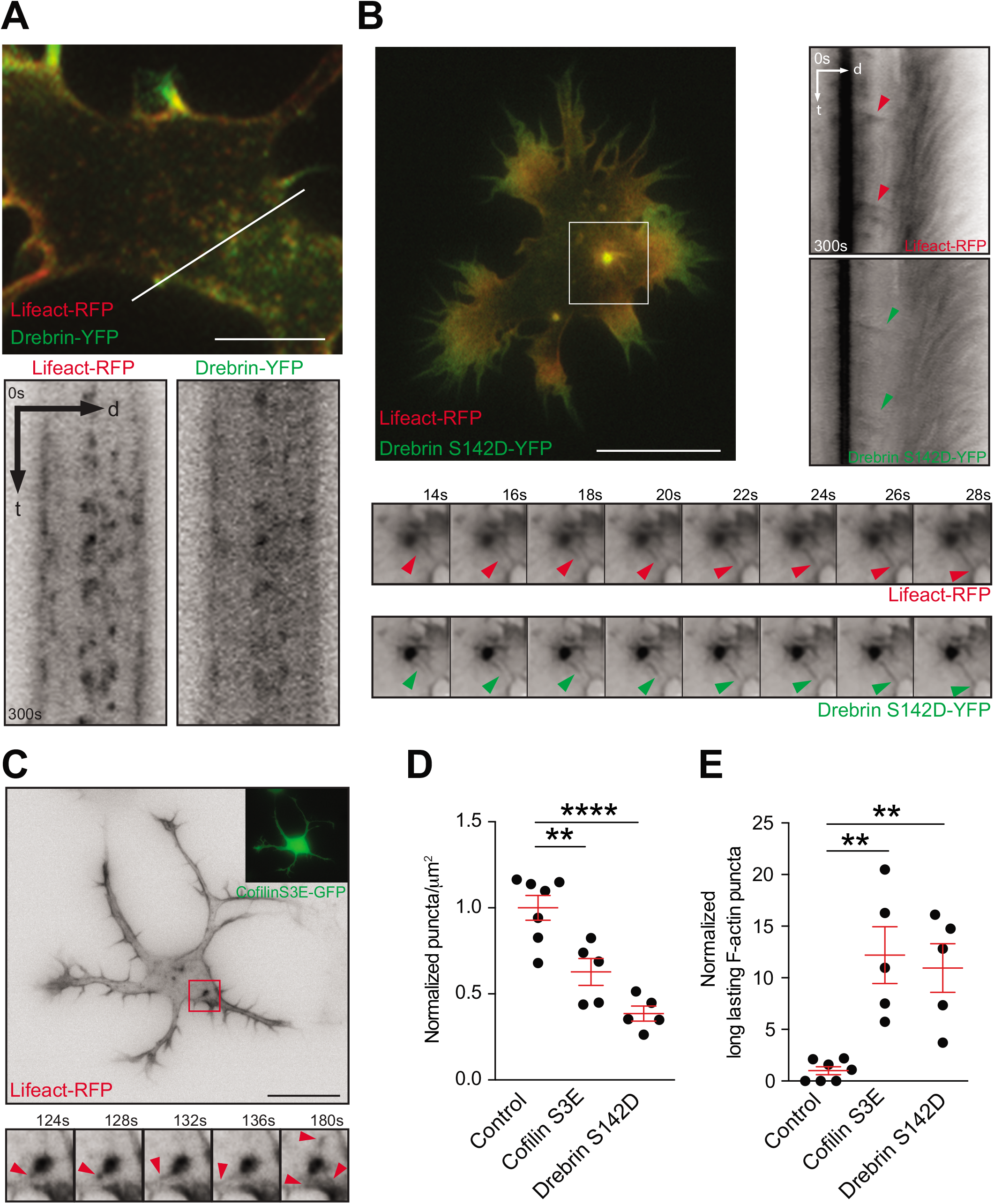
Stabilization of F-actin, by overexpressing Drebrin or Cofilin phosphomimetic mutant, unveils F-actin release from the puncta to the periphery. **(A)** Stage 2 cell transfected with Lifeact-RFP and Drebrin-YFP shows similar distribution of F-actin and Drebrin, highlighted via Kymograph. **(B, C)** Cells co-transfected with Lifeact-RFP and **(B)** Drebrin-S142D-YFP or **(C)** Cofilin-S3E-GFP is. Kymograph (in B) and time-lapse montages (in B and C), from insets, show F-actin asters releasing fibers to the cell periphery (arrow heads). **(D)** Density of somatic F-actin puncta of stage 2 cells in control group = 1.000 ± 0.07143, Cofilin-S3E group = 0.6277 ± 0.07836 and Drebrin-S142D group = 0.3856 ± 0.04340; p<0.0001 by one-way ANOVA, post hoc Dunnet test, **p<0.01, ****p<0.0001. **(E)** Comparison of long-lasting F-actin puncta in the soma of stage 2 cells. control group = 1.000 ± 0.3787, Cofilin-S3E group = 12.20 ± 2.746, Drebrin-S142D group = 10.95 ± 2.346; p=0.0009 by one-way ANOVA, post hoc Dunnet test, **p<0.01. Mean ± SEM; n=7 cells for control, n=5 cells each for Cofilin-S3E and Drebrin-S142D groups from at least two different cultures (for data shown in **D** and **E**). Scale bar: 5 μm (A) and 10 μm (B, C).

Likewise, expression of Cofilin-S3E decreased total number of somatic F-actin puncta with an increment of long-lasting F-actin puncta, compared to cells expressing only Lifeact-RFP (Fig. 5D, E), and reduced the F-actin treadmilling in growth cones (Fig. S11A). Furthermore, somatic F-actin puncta acquired an aster-like appearance releasing F-actin towards the cell cortex (Fig. 5C). Additionally, somatic F-actin fibers formed projections towards the cell periphery (Fig. S11B) Interestingly, F-actin travels along those F-actin fibers to reach the cell periphery concomitant with lamellipodia formation (Fig. S11B; Movie S8). Altogether, these results demonstrate that somatic F-actin puncta release F-actin towards the cell periphery.

### Centrosomal activities affect F-actin in the cell periphery

Given that somatic F-actin puncta concentrate near the centrosome (Fig. 1B), we asked whether centrosomal integrity is required for F-actin dynamics in developing neurons. We used chromophore-assisted light inactivation (CALI) based on the genetically encoded photosensitizer KillerRed, which upon green light illumination (540-580 nm), will specifically inactivate the target protein via the generation of light-activated reactive oxygen species (Bulina et al., 2006). We fused Centrin2, a protein confined to the distal lumen of centrioles and present in the pericentriolar material, to KillerRed (Centrin2-KR) to specifically inactivate the centrosome with laser irradiation (561 nm). Cells expressing Centrin2-KR and either EB3-GFP or Lifeact-GFP were imaged before laser irradiation. We then locally irradiated the centrosome with the 561nm laser for 1.5 sec to inactivate Centrin2 specifically at the centrosomal region without affecting somatic Centrin2 (Fig. 6A, B and Fig. S12A, B; Movie S9 and Movie S10). Two to three hours after laser irradiation, cells were reimaged. Centrosome inactivation via CALI reduced the number of somatic microtubules (Fig. S12A, B; Movie S9). Most importantly, F-actin speed treadmilling as well as the F-actin intensity at the cell periphery were significantly reduced after centrosomal disruption (Fig. 6A, B; Movie S10). As a control we irradiated a similar sized area at the soma away from the centrosome with the same settings. The cells irradiated outside the centrosomal area did neither show reduced F-actin treadmilling speed nor decreased F-actin intensity in growth cones (Fig. S12C, D; Movie S11).

**Fig. 6.**
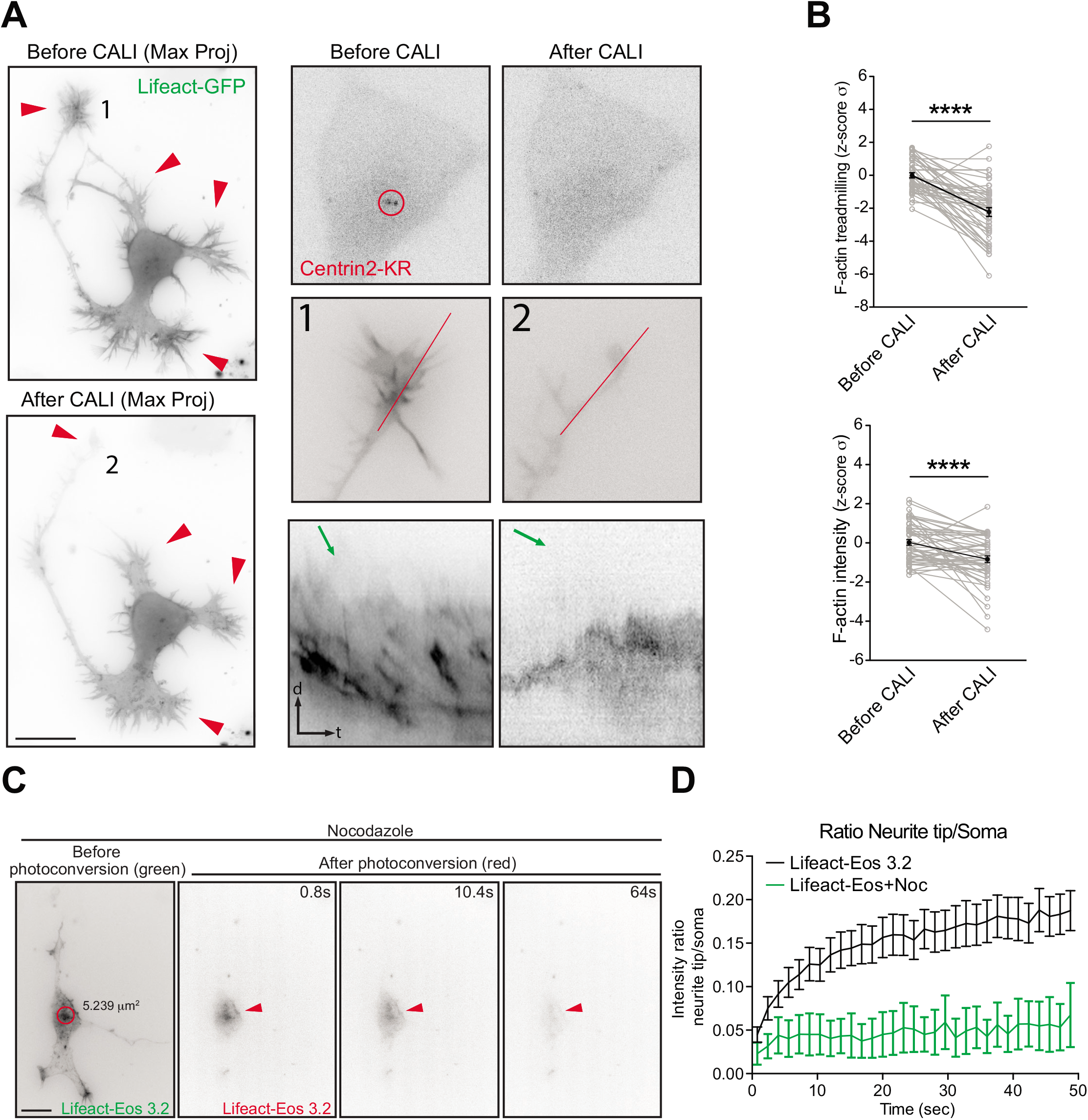
Centrosome inactivation affects F-actin intensity and treadmilling in growth cones. **(A)** Neuron transfected with Centrin2-KillerRed and Lifeact-GFP subjected to localized CALI treatment. Upper panel: Centrin2-KR signal before and after treatment; Middle and lower panels: Kymograph of actin treadmilling in growth cone before and after CALI treatment **(B)** Influence of CALI on F-actin treadmilling speed and F-actin intensity in growth cones. Mean ± SEM; n=10 cells from at least three different cultures. **(C)** Nocodazole (7uM) treated Lifeact-mEos3.2 expressing neuron photoconverted in the soma with 405 nm laser (red circle). Cell before (green) and after (red) photoconversion. Arrowheads point photoconverted signal movement. **(D)** Growth cone to soma photoconverted signal intensity ratios of untreated and Nocodazole treated Lifeact-mEos3.2 expressing cells. Experiments shown in Figure 6 C, D, Supplementary figure 8 D, E and Supplementary figure 10 C, D were done at the same time, therefore the same Control data is used for comparison. Mean ± SEM; n=12 cells for untreated and n=8 cells for nocodazole groups from at least two different cultures. Scale bar: 10 μm (A, C).

Microtubule organization in early developing neurons is centrosome-dependent (Calderon de Anda et al., 2008; Stiess et al., 2010; Yau et al., 2014). Therefore, we decided to disrupt microtubule polymerization with nocodazole to test whether the F-actin translocation towards the cell periphery is affected. We found that nocodazole drastically reduced the motility of somatic photoconverted Lifeact-mEos3.2 (Fig. 6C, D; Movie S12). Accordingly, microtubules disruption in developing neurons leads to a less dynamic F-actin cytoskeleton (Zhao et al., 2017). Altogether, these results show that the centrosome and microtubules are necessary for somatic F-actin translocation towards the cell periphery.

### PCM-1 determines somatic F-actin organization

Next, we tested PCM-1 as a molecular determinant of F-actin dynamics near the centrosome. PCM-1 promotes F-actin polymerization in non neuronal cells (Farina et al., 2016). We found that PCM-1 particles intermingled with F-actin puncta in the soma of fixed neurons and concentrated in proximity of F-actin puncta (Average distance between F-actin puncta-PCM-1 = 0.584 ± 0.019 μm; Fig. 5A, B). Accordingly, neurons transfected with PCM-1-GFP showed PCM-1-GFP granules surrounding and “touching” somatic F-actin puncta (Fig. S13A; Movie S13).

To further test whether PCM-1 and somatic F-actin organization are interrelated, we treated neurons (24 hrs after plating) with Cytochalasin D or Jasplakinolide. We found that polarized F-actin structures induced by Cytochalasin D or Jasplakinolide treatment (Fig. S8 A, B and Fig. S10 A, B) are accompanied by PCM-1 particles (Fig. S13B, C). Interestingly, when cells were co-treated with Cytochalasin D and Nocodazole, disperse F-actin clusters (96.97%, 66 cells from at least three different cultures) associated with PCM-1 particles formed (Fig. S13D). These data indicate that somatic F-actin organization is linked to PCM-1 and microtubules integrity.

To probe the involvement of PCM-1 more specifically we took advantage of *in utero* electroporation to introduce a PCM-1 shRNA construct to silence PCM-1 expression in cortical neurons and neuronal progenitors ((de Anda et al., 2010; Ge et al., 2010); Fig. S14A, C). We tested the role of PCM-1 in F-actin dynamics and neurite outgrowth of cultured developing neurons and neurons differentiating in the developing cortex. PCM-1 down-regulation in cultured neurons led to the formation of long and thin neurites (Fig. 7C; Fig. S14A, B, C and D), similar to the well-known effect induced by pharmacological F-actin disruption using cytochalasin D (Bradke and Dotti, 1999). This suggests that PCM-1 down-regulation impaired F-actin dynamics and thus boosted neurite outgrowth.

**Fig. 7.**
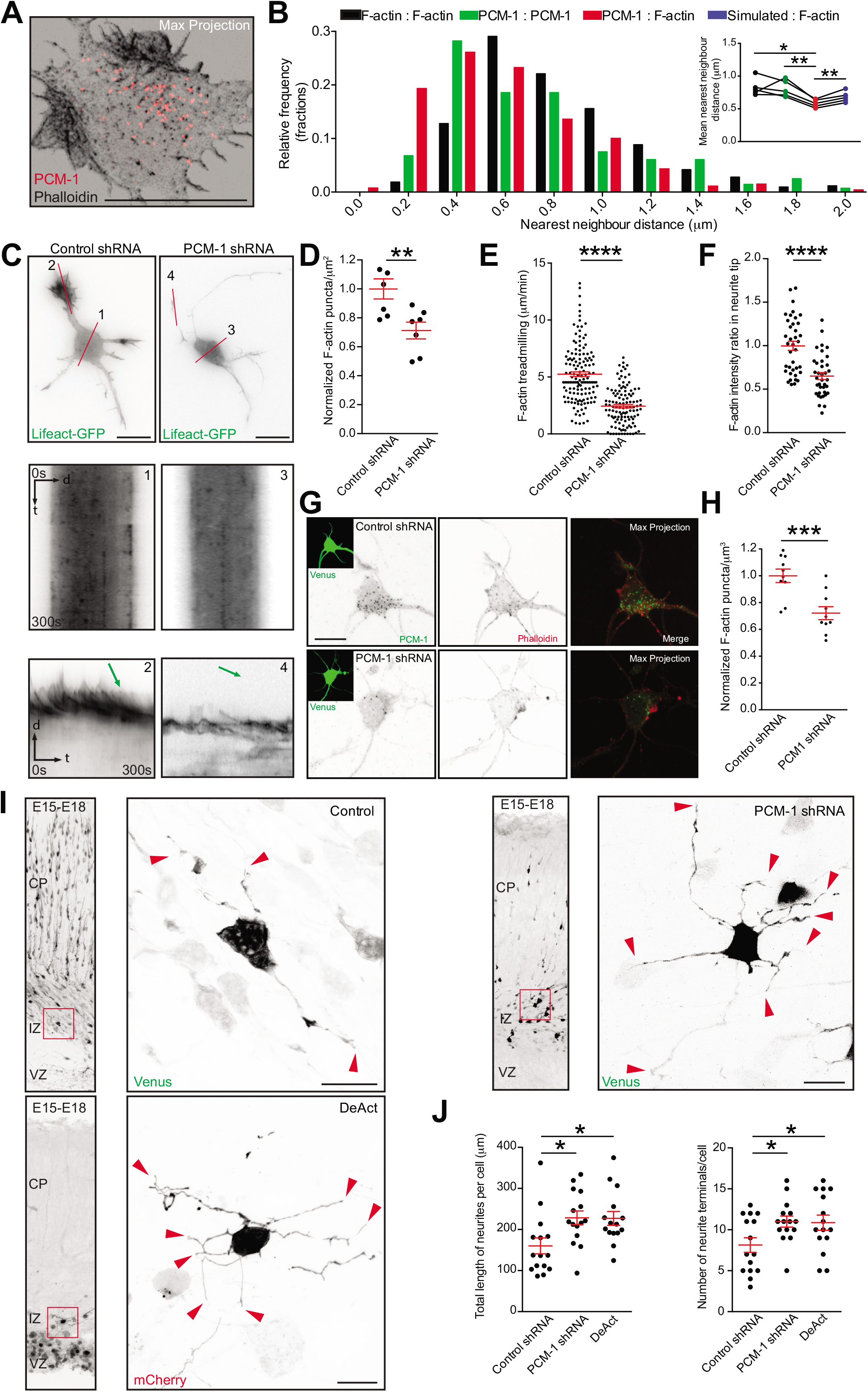
PCM-1 intermingles with F-actin puncta and is essential for the maintenance of F-actin in the soma and growth cones. **(A)** Confocal Max projection of a stage 2 neuron stained with PCM-1 antibody and phalloidin showing polarized and intermingled PCM-1 and F-actin puncta. **(B)** Nearest neighbor analysis showing frequency distribution of distance between F-actin: F-actin puncta (black bars), PCM-1: PCM-1 puncta (green bars) and PCM-1: F-actin puncta (red bars). Inset: paired mean nearest neighbor distance values of F-actin:F-actin (0.827 ± 0.058), PCM-1:PCM-1 (0.811 ± 0.057), PCM-1:F-actin (0.584 ± 0.026) and simulated:F-actin (0.683 ± 0.036) puncta in the soma from n = 5 cells, Mean ± SEM. **(C)** DIV I cortical neurons transfected with either control or PCM-1 shRNA together with Lifeact-GFP. Kymographs are obtained from lines marked as 1 and 3 for soma of control and PCM-1 shRNA, respectively. Lines 2 and 4 for growth cones of control and PCM-1 shRNA, respectively. **(D)** Density of somatic F-actin puncta of stage 2 cells from control condition = 1.000 ± 0.0690, PCM-1 shRNA condition = 0.7122 ± 0.05809; **p= 0.0078 by unpaired Student’s t-test. Mean ± SEM; n=7 cells each for control and PCM-1 shRNA from at least three different cultures. (E) F-actin treadmilling speed (μm/min) in growth cones (or neurite tips from PCM-1 shRNA-expressing cells) of stage 2 cells from control condition = 5.2452 ± 0.2064; PCM-1 shRNA condition = 2.4402 ± 0.1543; ****p<0.0001 by unpaired Student’s t-test. Mean ± SEM; n=10 cells for control and PCM-1 shRNA from at least three different cultures. **(F)** F-actin intensity ratio in growth cones of control condition = 0.9979 ± 0.0526, neurite tips of PCM-1 shRNA condition = 0.6513 ± 0.0401. ****p<0.0001 by unpaired Student’s t-test. Mean ± SEM; n=10 cells for control, n = 8 cells for PCM-1 shRNA groups from at least two different cultures. **(G, H)** PCM-1 down-regulation decreased the density of somatic F-actin puncta detected with phalloidin staining of stage 2 cells from control condition = 1.000 ± 0.0500, PCM-1 shRNA condition = 0.7212 ± 0.0485; p= 0.0008 by unpaired Student’s t test. Mean ± SEM; n = 10 cells each for control and PCM-1 shRNA groups from at least three different cultures. **(I)** PCM-1 downregulation or actin depolymerization (via DeAct expression) result in neurite elongation and increase in number of neurite terminals *in vivo*. Inset: Multipolar cells in the intermediate zone (IZ) expressing control shRNA, PCM-1 shRNA or DeAct plasmids together with Venus or mCherry. **(J)** Left panel: Total length of neurites per cell in control condition = 160 ± 19.37, PCM-1 shRNA = 228.5 ± 16.82, and DeAct expressing cells = 227 ± 16.71. p=0.0123 by oneway ANOVA, post hoc Dunnett’s test. *p<0.05. Right panel: Number of neurite terminals per cell in control condition = 8.133 ± 0.88, PCM-1 shRNA = 11 ± 0.66, and DeAct expressing cells = 10.87 ± 0.93. p=0.0321 by one-way ANOVA, post hoc Dunnett’s test. *p<0.05. Mean ± SEM; n = 15 cells from three brains in each group. Scale bar: 10 μm (A, C, G and I).

In extension of these findings, we observed a direct effect of PCM-1 down-regulation on F-actin dynamics with a reduced total number of F-actin puncta in the cell body detected with Lifeact-GFP (Fig. 7C, D) or Phalloidin (Fig. 7G, H). Moreover, using specific actin nucleator inhibitors (SMIFH2 and CK666), we were able to show that the somatic F-actin puncta are Formin-but not Arp2/3-dependent (Fig. S15). Furthermore, PCM-1 down-regulation significantly decreased the F-actin treadmilling speed (Fig. 7C, E) as well as the relative F-actin levels in neurite tips (Fig. 7C; F). Of note, the effects of PCM-1 knockdown were reversed when an RNAi resistant plasmid, Chicken-PCM-1-GFP, was transfected along with Lifeact-RFP and PCM-1-shRNA (Fig. S16).

Finally, we tested whether PCM-1 down-regulation or F-actin disruption similarly affect neuronal differentiation in the developing cortex. We electroporated *in utero* control shRNA or PCM-1 shRNA, together with Venus as well as DeAct plasmid, which impairs F-actin dynamics (Harterink et al., 2017), at E15 to analyze the neuronal morphology at E18 *in situ*. Importantly, we found that down-regulating PCM-1 in the developing cortex and disrupting F-actin in newly born neurons promotes neurite elongation in a similar manner (Fig. 7I, J). Thus, suggesting that PCM-1 down-regulation affects the amount of somatic F-actin, which is produced to modulate neurite outgrowth. Altogether our results show that PCM-1 regulates somatic F-actin dynamics and that somatic actin polymerization has an effect on growth cone dynamics.

## Discussion

Collectively, our results indicate that i) F-actin in the cell body organizes around the centrosome and ii) somatic F-actin is released towards the cell periphery, thus affecting growth cone behavior. To our knowledge, the neuronal F-actin organization described here is a novel cellular mechanism to sustain neuronal development. Although our data do not clarify the molecular mechanism by which somatic F-actin is delivered towards the cell periphery, our results suggest that microtubule organization is relevant for somatic F-actin delivery to growth cones.

Mechanistically, we show that this somatic F-actin organization in neurons relies on the presence of PCM-1. PCM-1 intermingles with somatic F-actin aster-like structures, which concentrate near the centrosome. Our time-lapse analysis further corroborates this PCM-1/somatic F-actin organization. Finally, and most importantly, our data show that PCM-1 down-regulation affects F-actin dynamics in neurite tips of developing neurons and hence neuronal differentiation *in vitro* and *in vivo*. To our knowledge, this is the first time PCM-1 has been associated with F-actin dynamics in neurons and we are the first to show that PCM-1-dependent somatic F-actin organization has a direct effect on distal growth cones behavior.

However, our data do not show how PCM-1 regulates the somatic F-actin polymerization. Farina et al., (2016) described that centrosomal F-actin polymerization is Arp2/3 dependent. In contrast, we found that formins but not Arp2/3 contribute to the somatic F-actin organization in neurons, as shown for axonal F-actin organization of mature neurons (Chakrabarty et al., 2018; Ganguly et al., 2015). Thus, the mechanism described by Farina et al. (2016) may not be the same in neurons. Further experiments need to be performed in neurons to ensure a deep understanding of the somatic F-actin nature described here. In summary, we believe our data will pave the way to future important contributions oriented to understand F-actin organization and dynamics in developing neurons.

## Supporting information

## Acknowledgements

Thanks to M. Kneussel, M. Harterink (C. Hoogenraad’s laboratory), T. Oertner and X. Chai (M. Frotscher’s labaratory) for plasmid constructs. Thanks to the laboratories of M. Kneussel and P. Soba at the ZMNH for equipment use. Thanks to I. Hermans-Borgmeyer and members of UKE-Hamburg animal facility for their help with animal experiments.

M. Kreutz is supported by the Leibniz Foundation, Deutsche Forschungsgemeinschaft (Kr1879 5-1, 6-1; SFB 779 TPB8), BMBF Energi and JPND STAD. M. Mikhaylova is supported by grants from the Deutsche Forschungsgemeinschaft (DFG Emmy-Noether Programm (Ml 1923/1-1) and FOR2419 (MI 1923/2-1). O. Kobler is funded by the DFG Grant SCHE 132/18-1. F. Calderon de Anda is supported by Deutsche Forschungsgemeinschaft (DFG) Grants: FOR 2419, CA1495/1-1, and CA 1495/4-1; ERA-NET Neuron Grant (Bundesministerium für Bildung und Forschung, BMBF, 01EW1410 ZMNH AN B1), Landesforschungsförderung Hamburg (Z-AN LF), and University Medical Center Hamburg-Eppendorf (UKE). DP. Meka is a co-applicant in the Deutsche Forschungsgemeinschaft (DFG) Grant CA 1495/4-1 for F. Calderon de Anda.

---

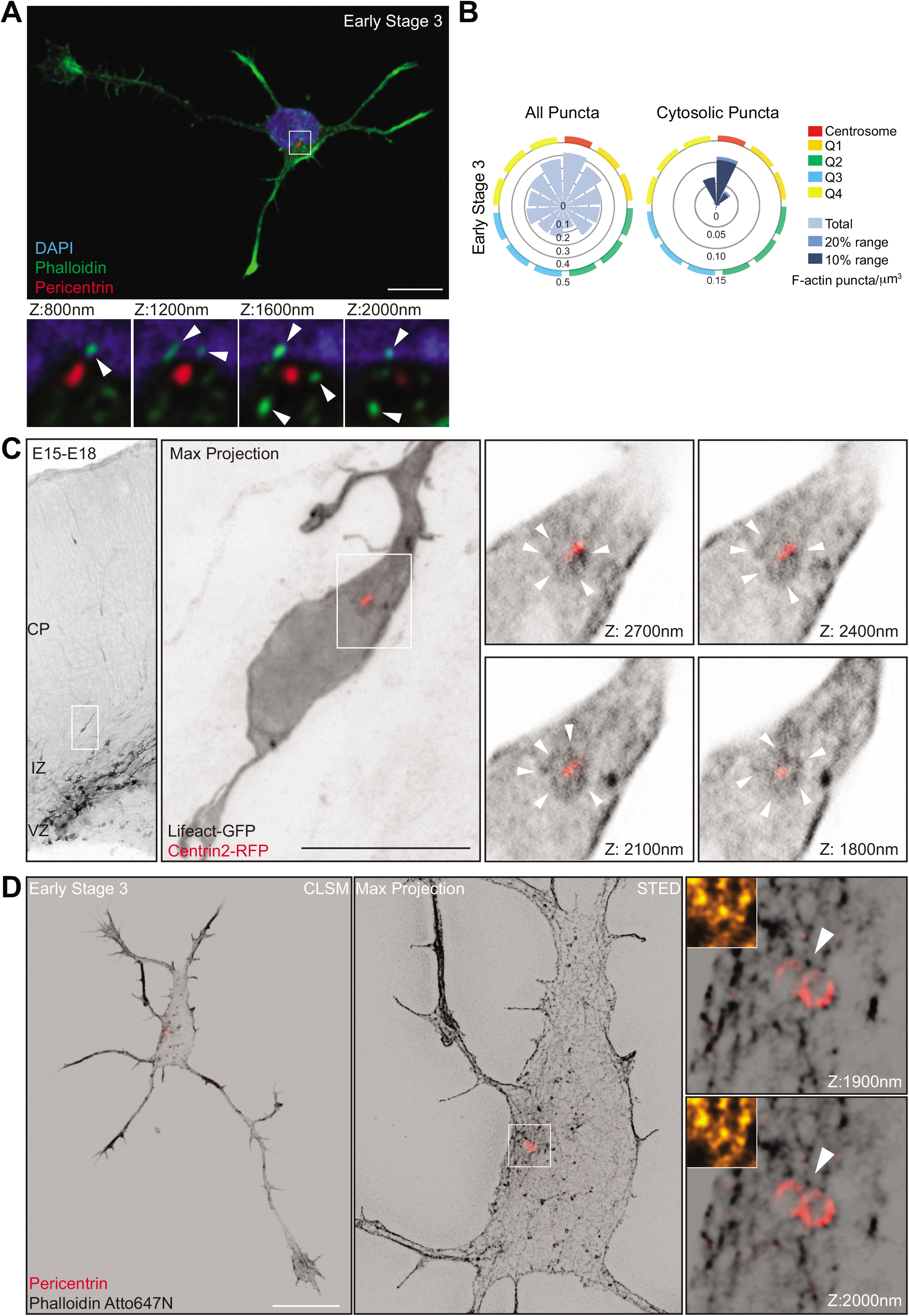

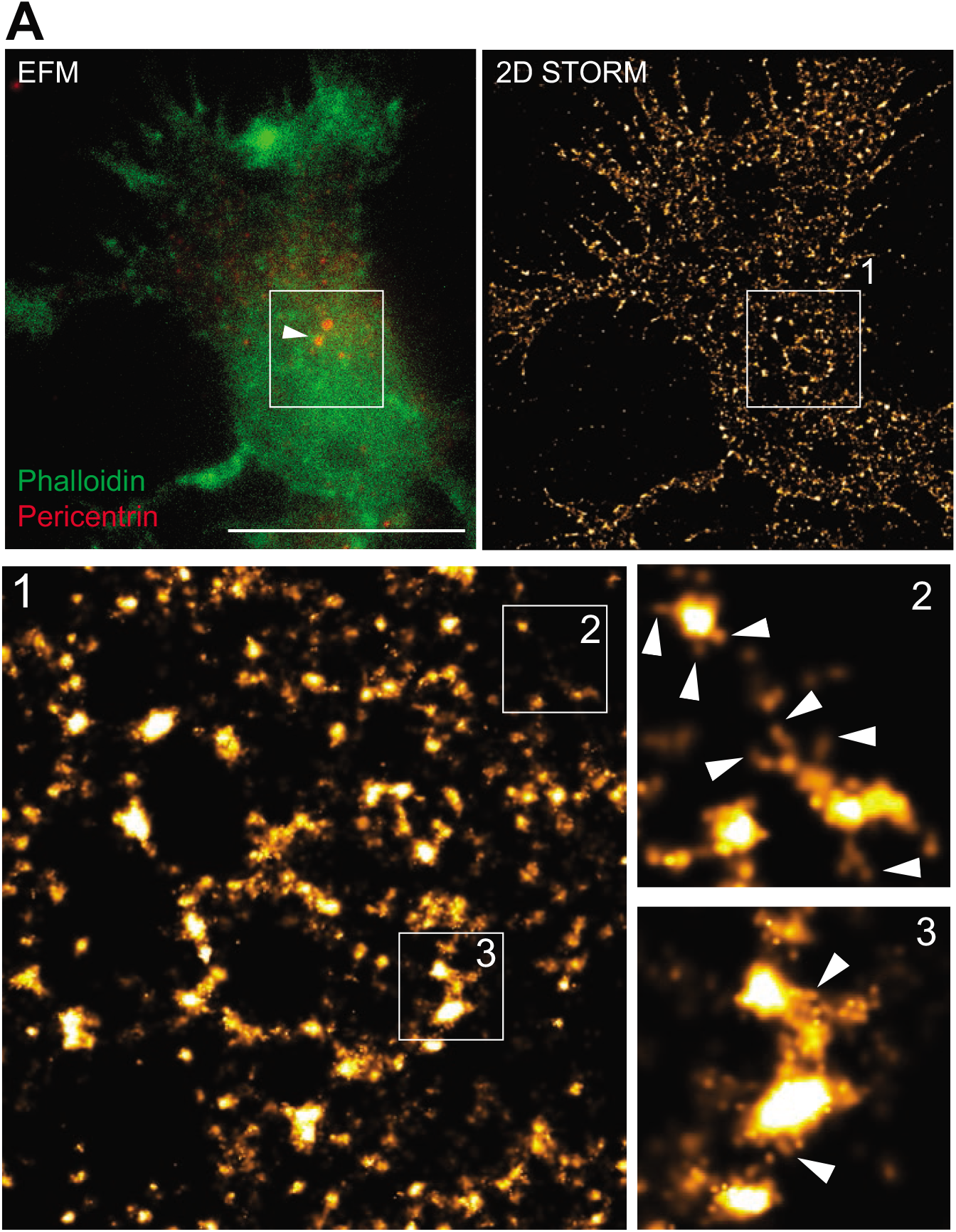

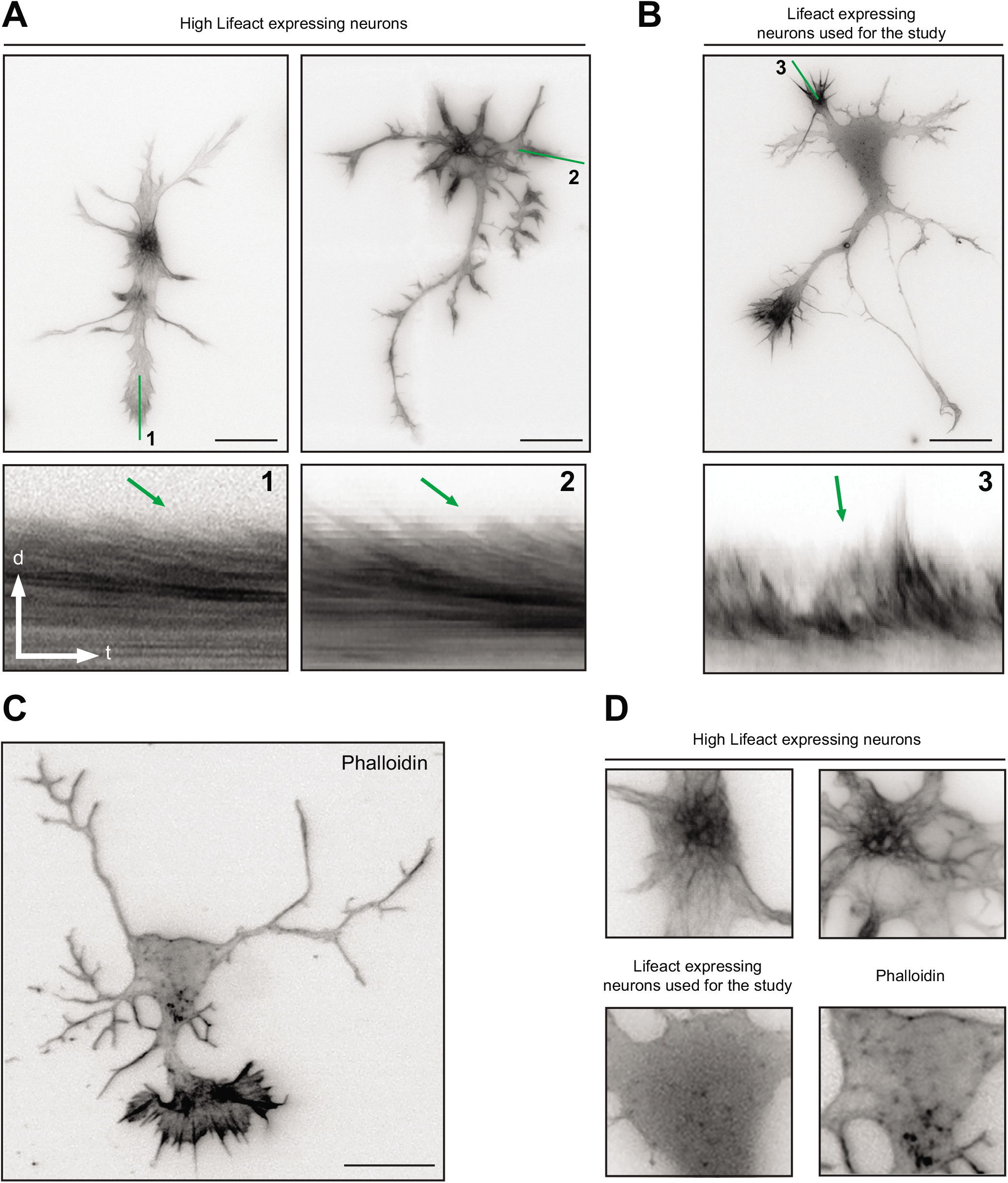

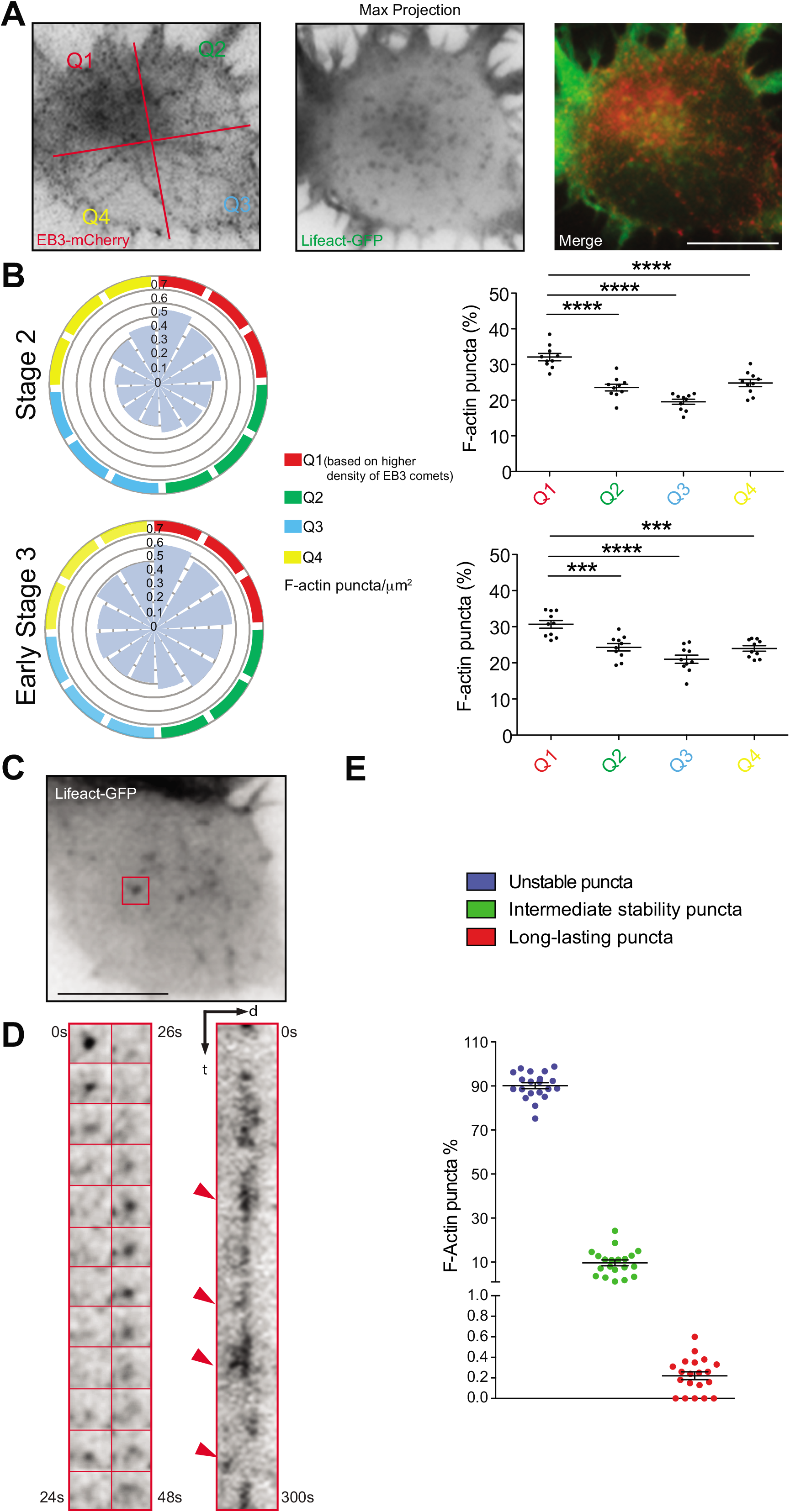

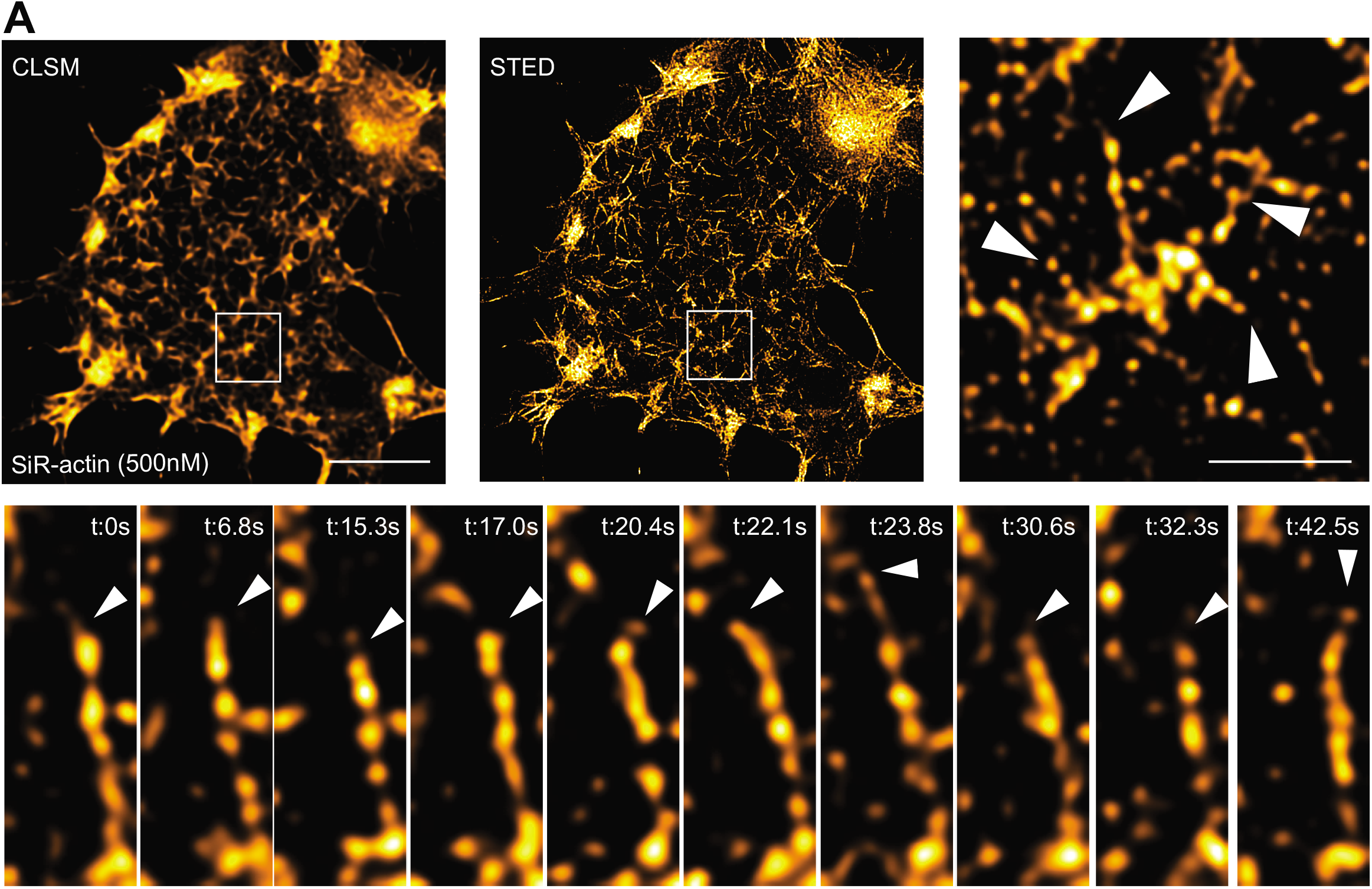

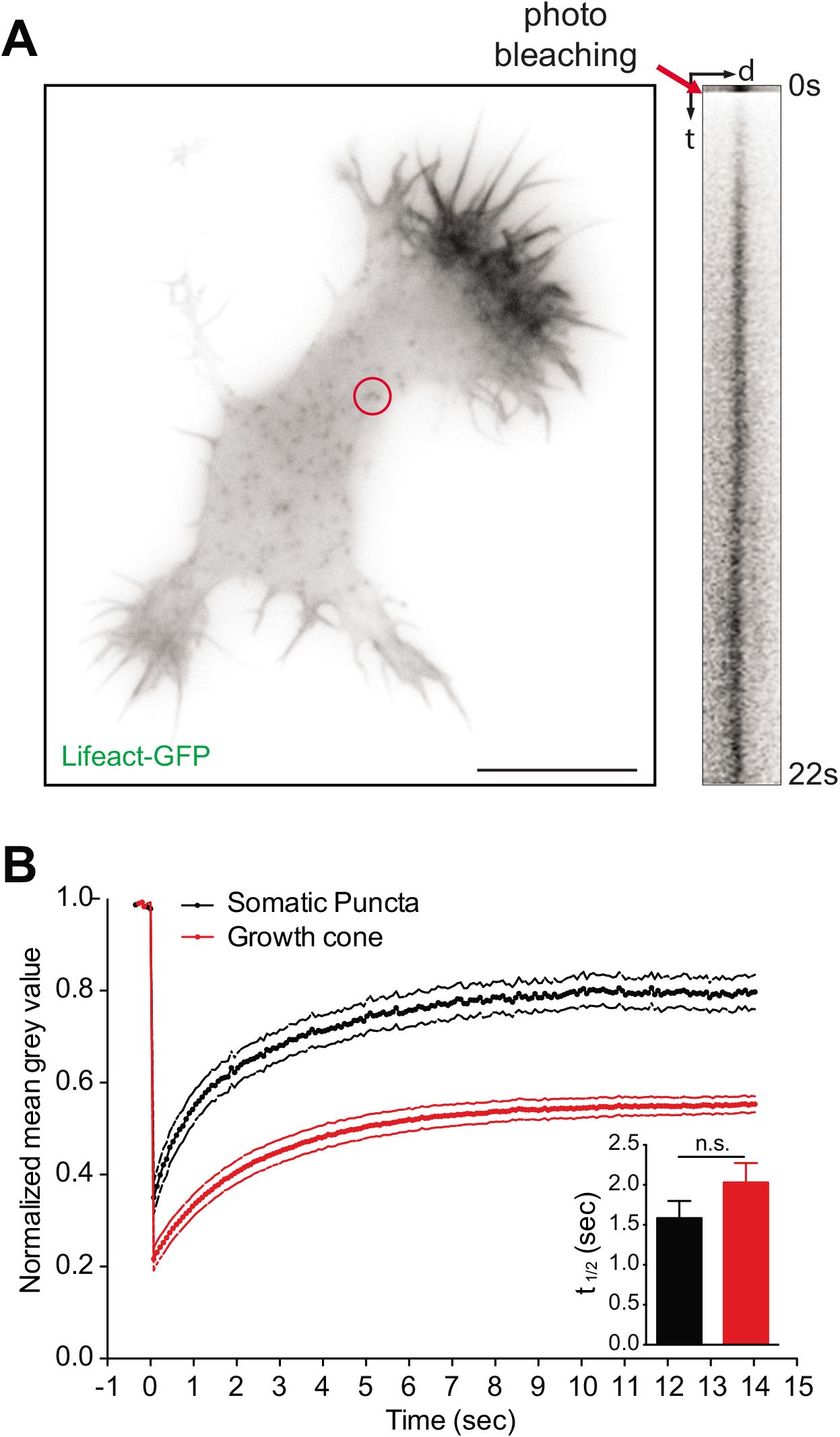

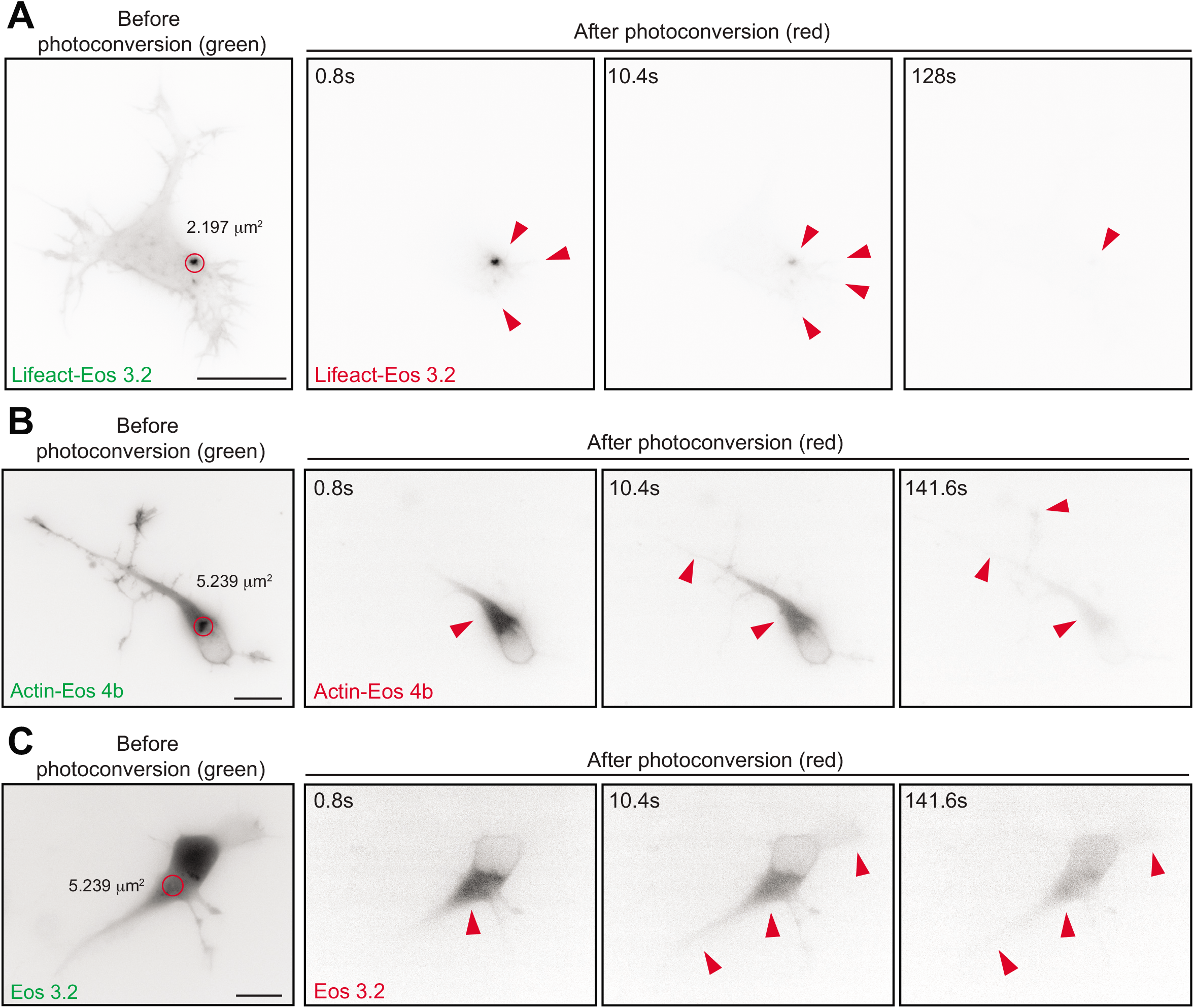

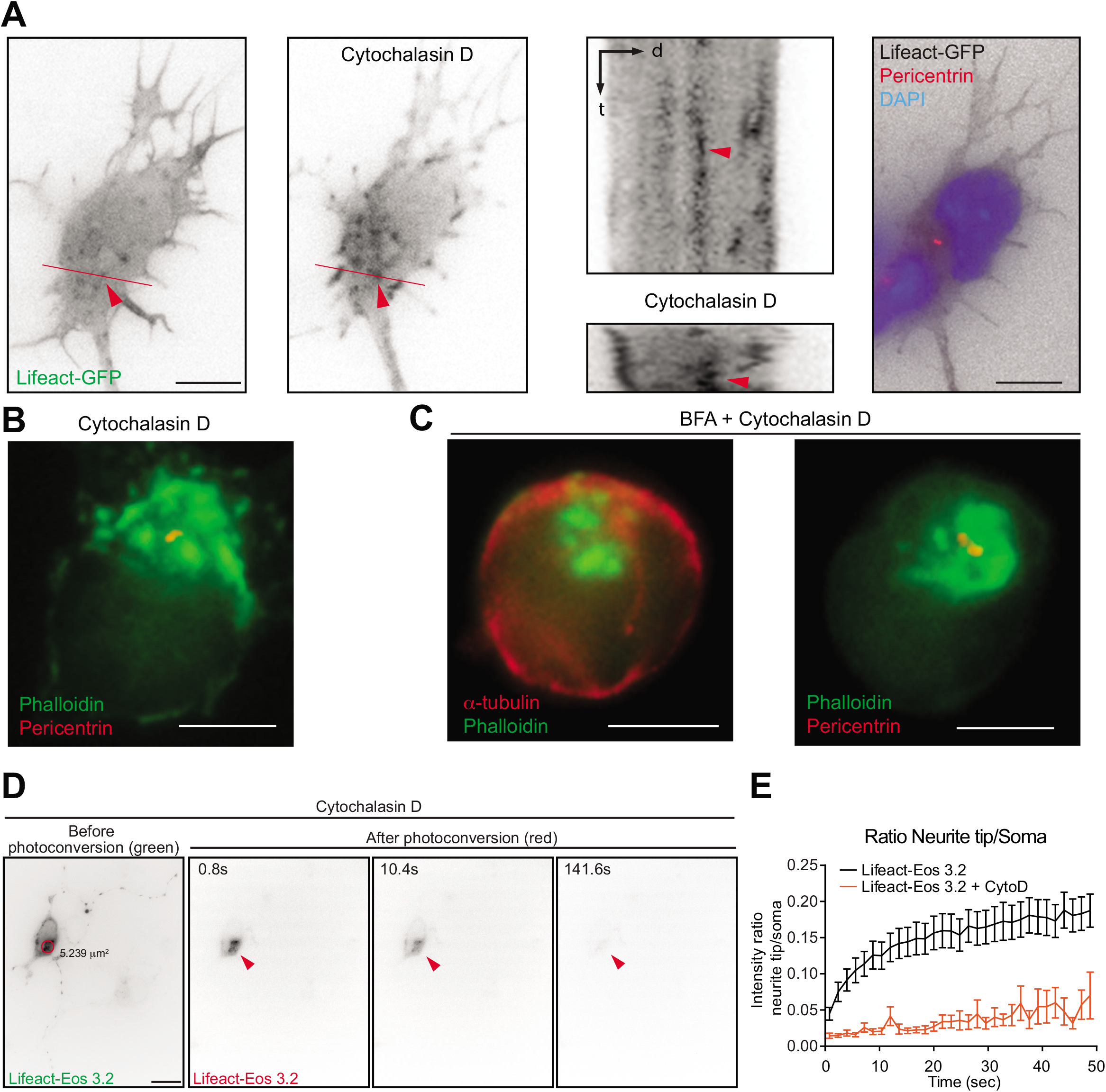

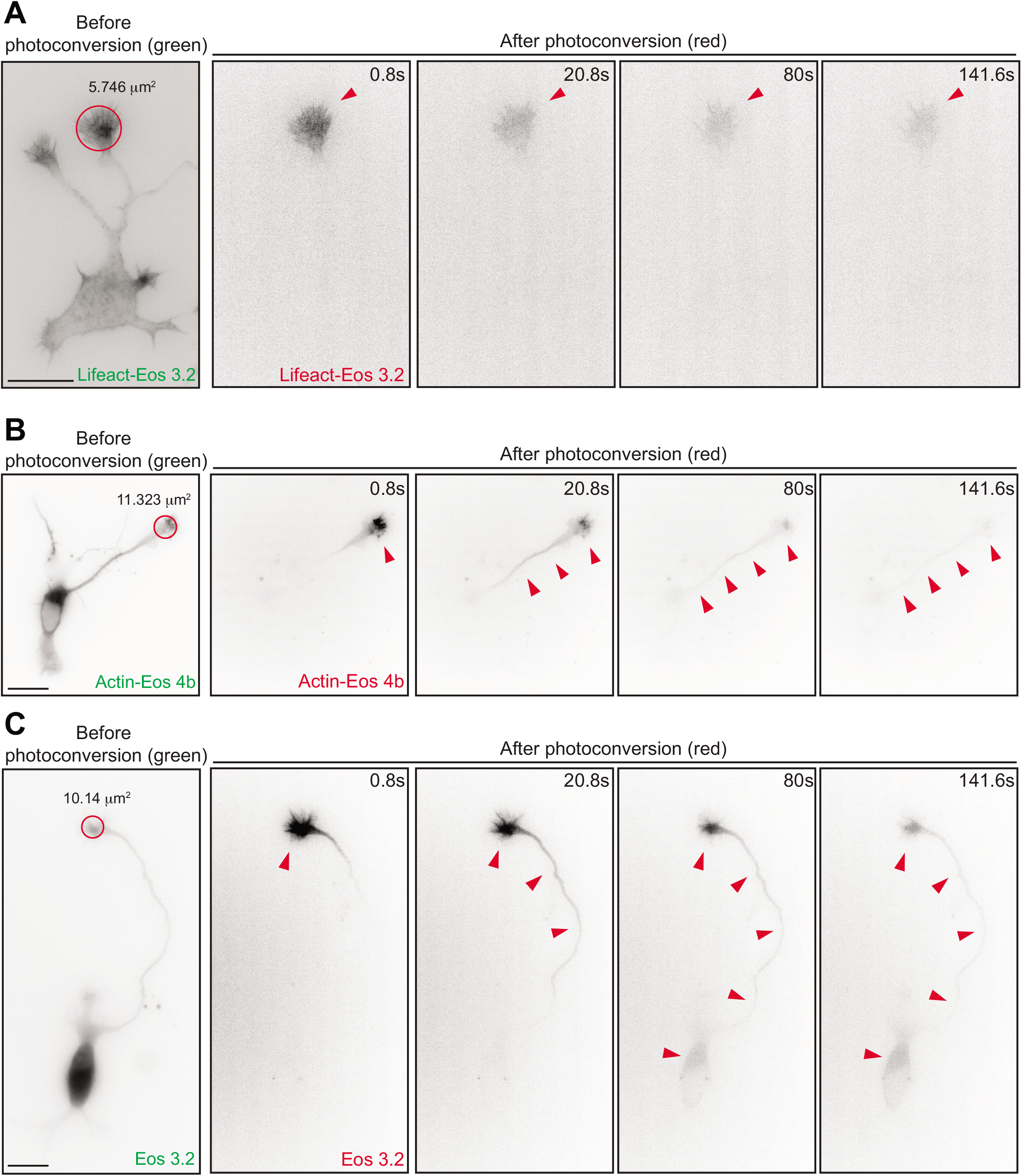

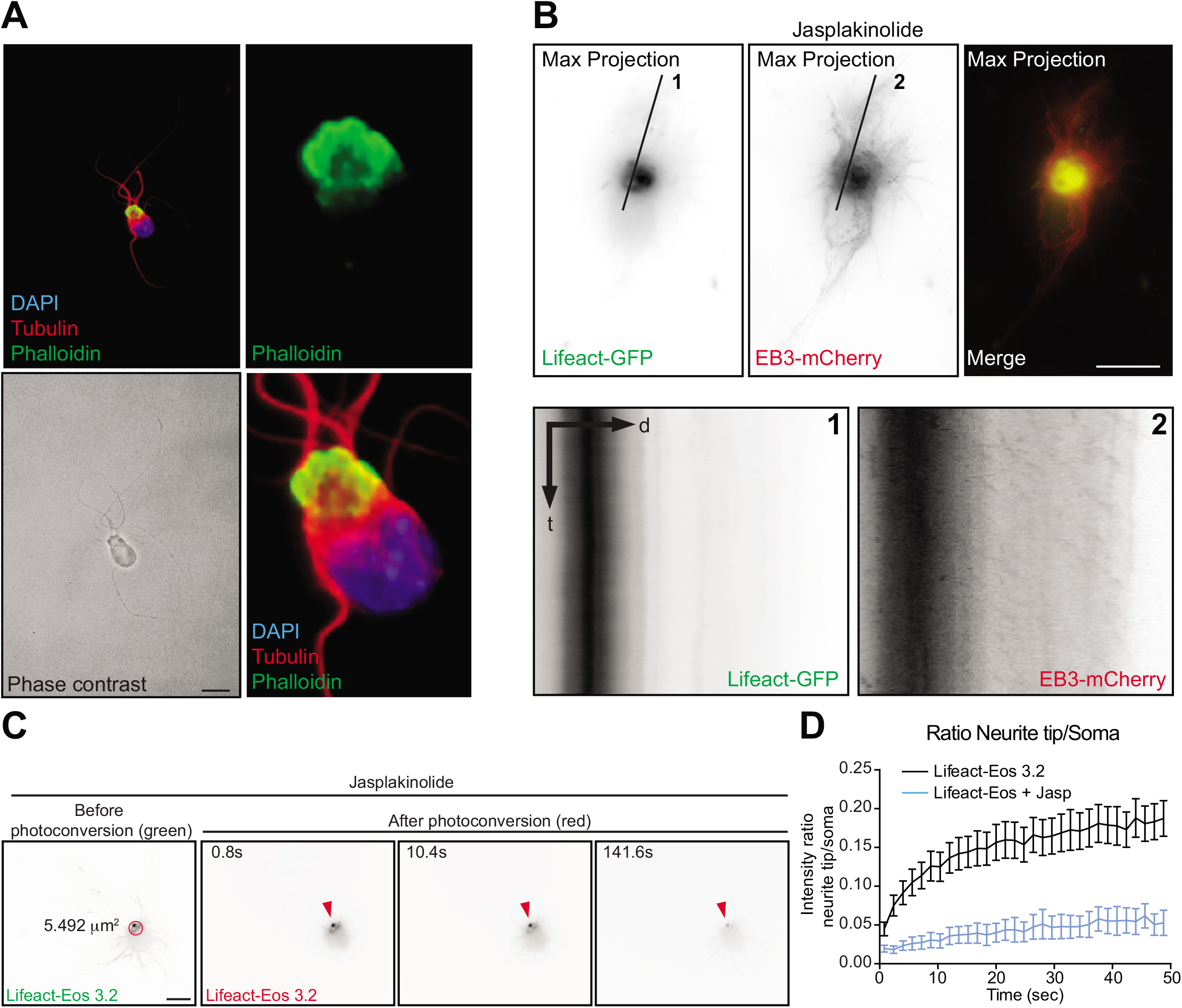

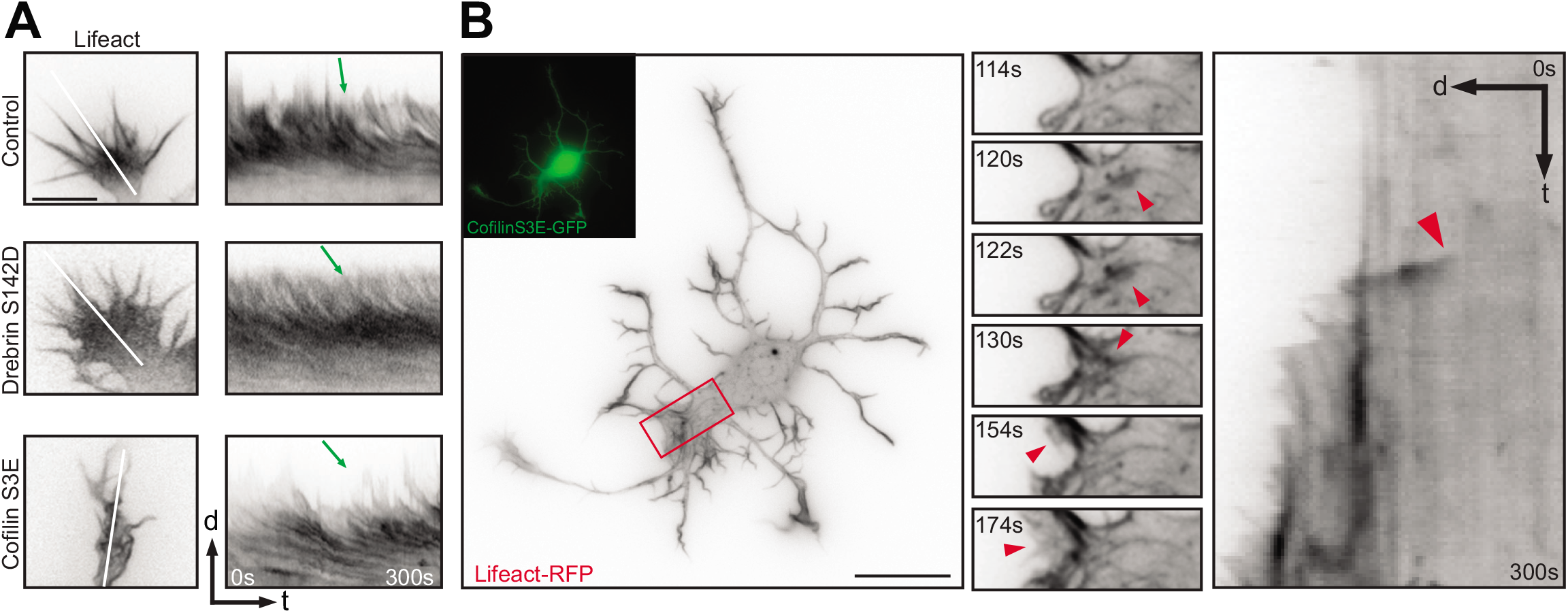

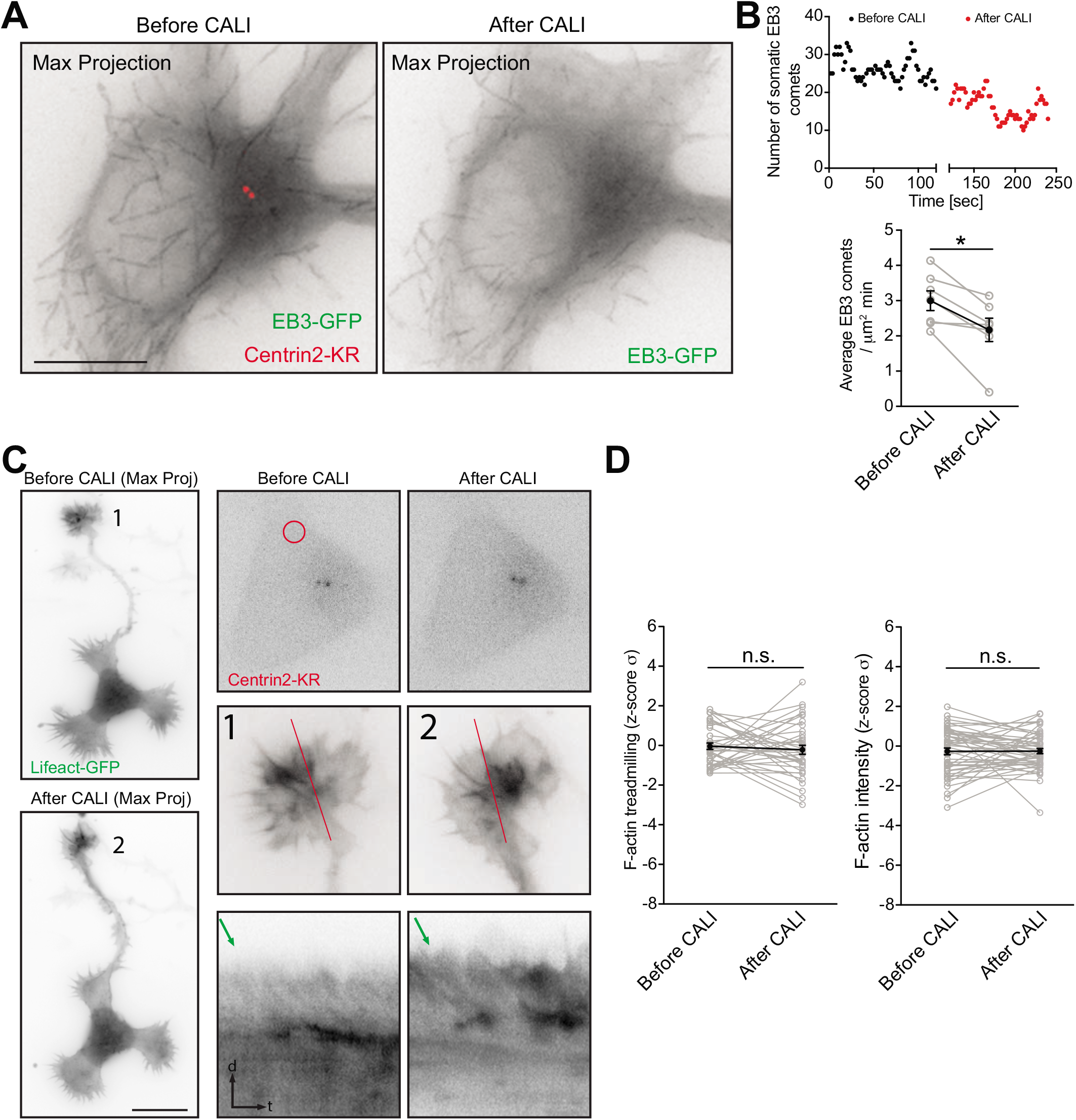

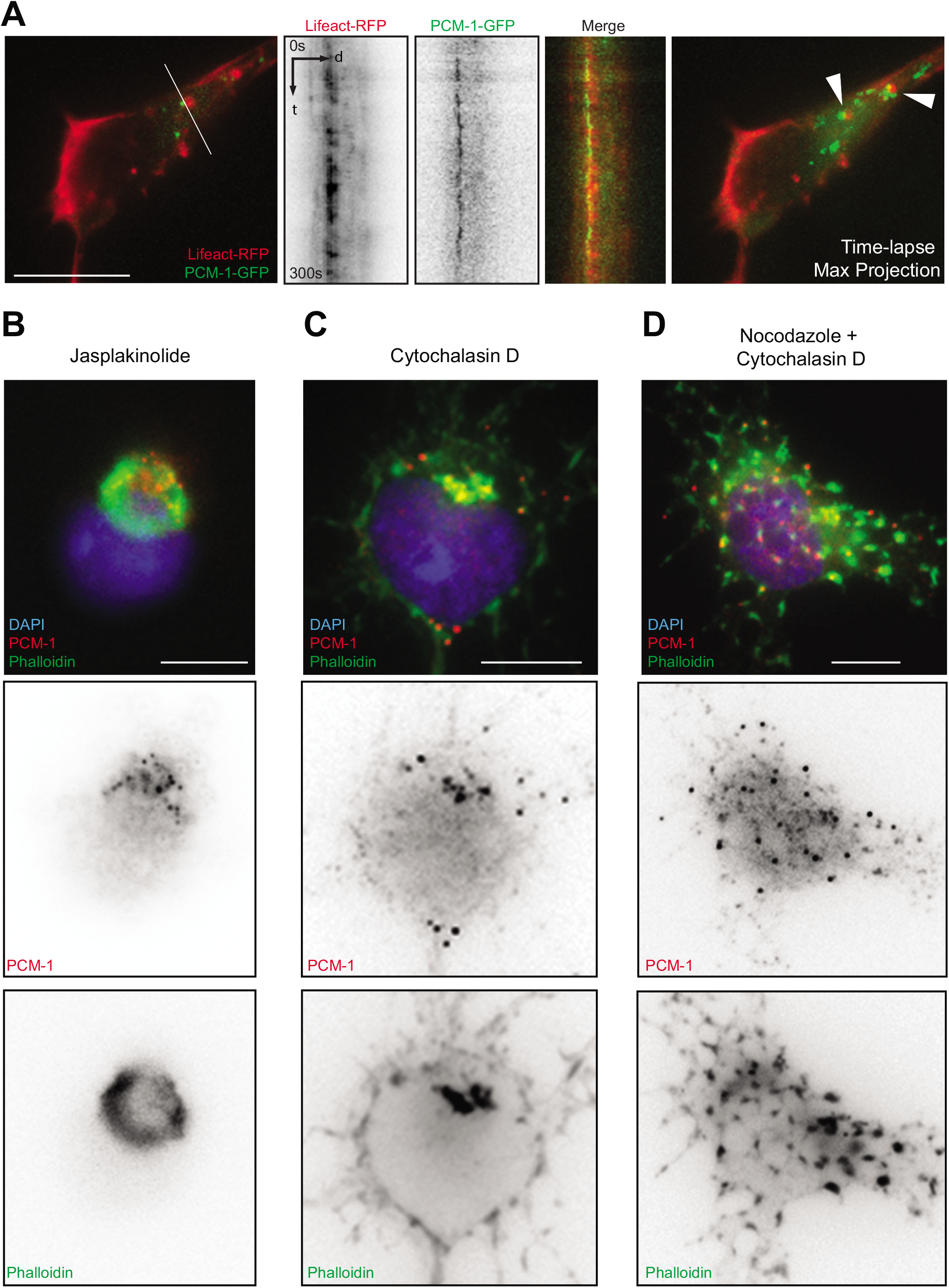

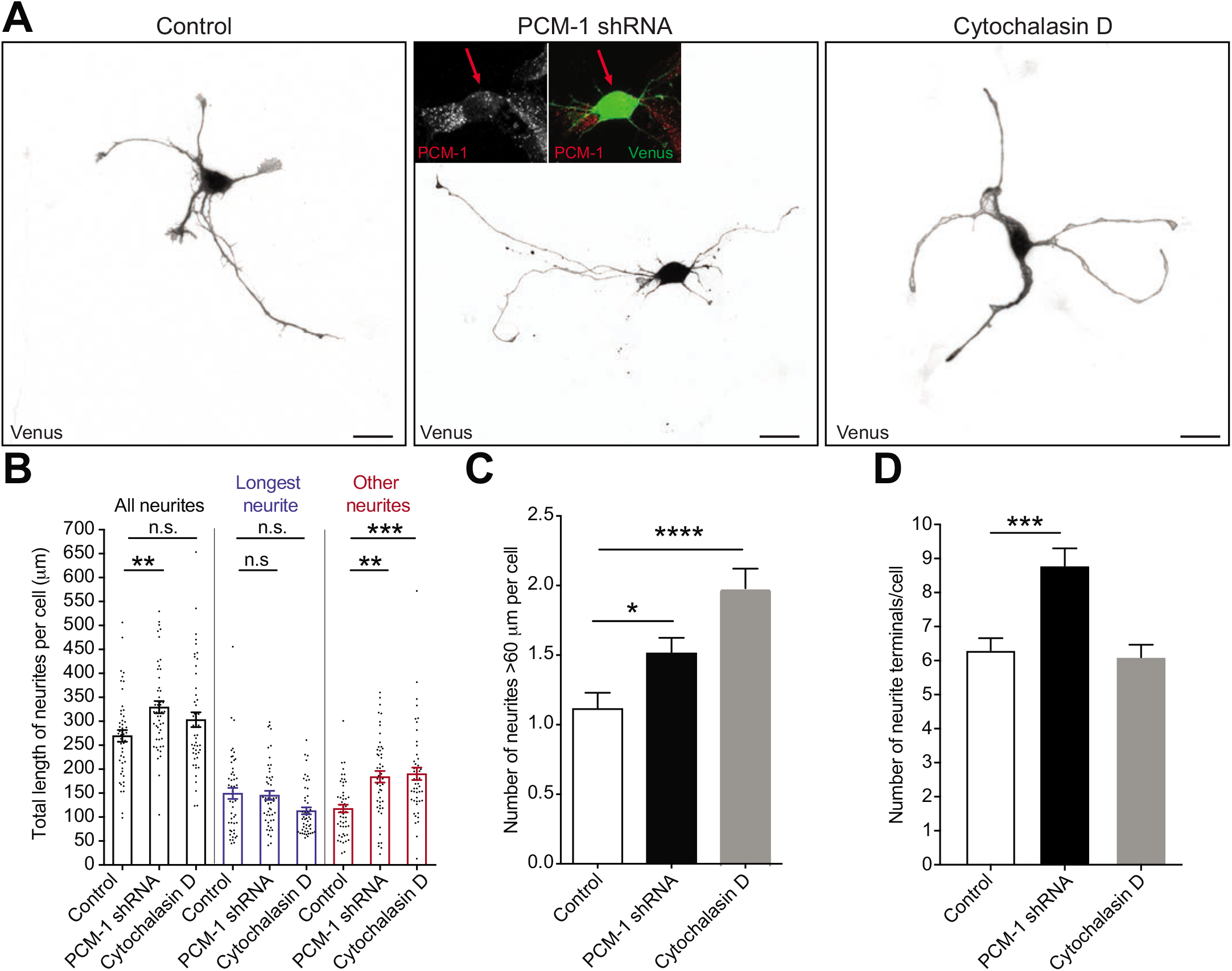

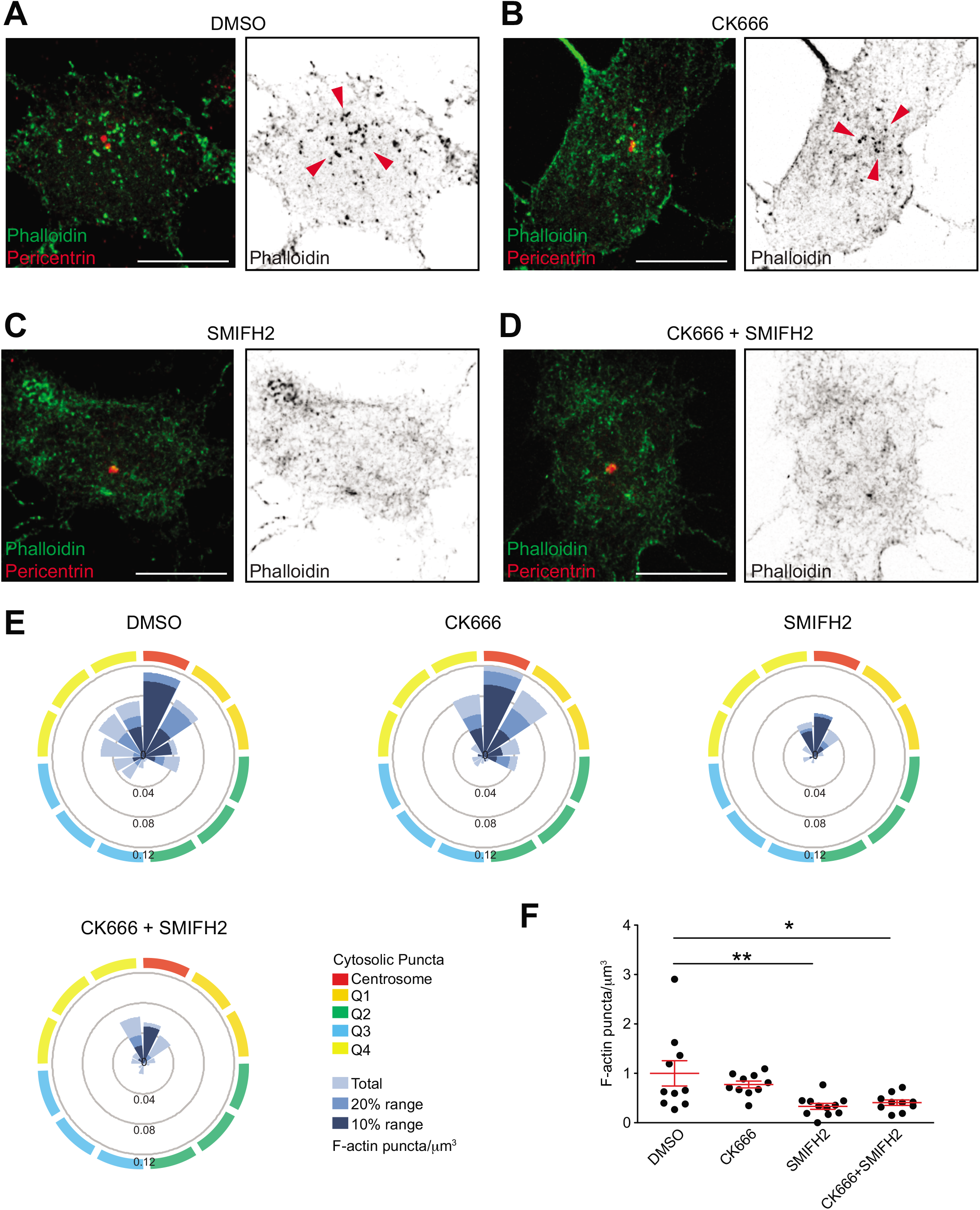

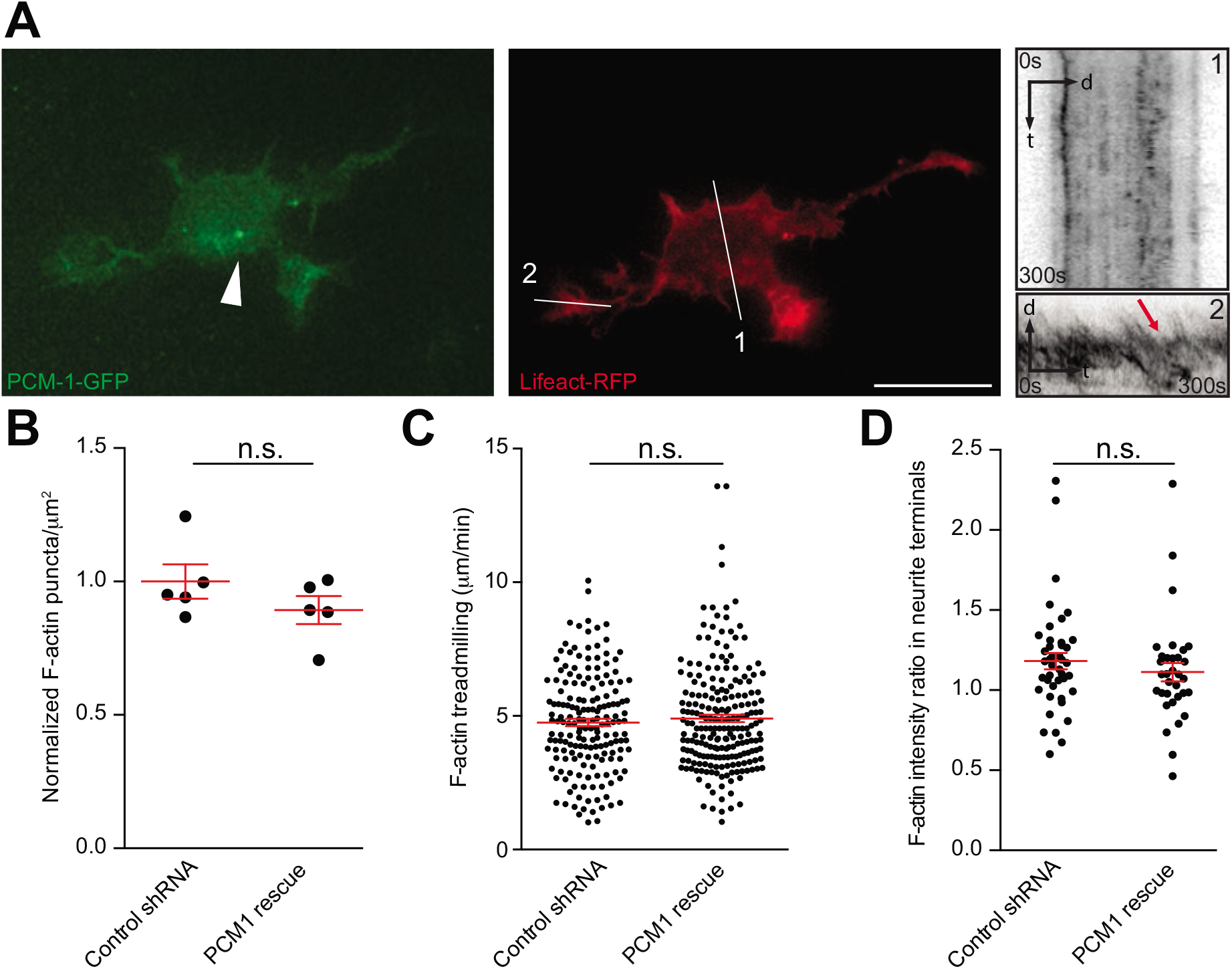

