## Supplementary material for "Radial F-actin Organization During Early Neuronal Development"

**Supplementary Materials.**

**Material and Methods.**

**RNAi and fluorescent protein constructs.**

We previously reported the Venus, mCherry, Control shRNA, PCM-1 shRNA and Chicken PCM-1-GFP plasmids (de Anda et al., 2010; Ge et al., 2010). A pSilencer vector containing a random sequence hairpin insert was used as a control for the shRNAs (de Anda et al., 2010; Ge et al., 2010). M. Harterink (from the laboratory of C. Hoogenraad, Utrecht University) kindly provided the DeAct-SpvB plasmid (that was cloned into the pGW2-GFP vector) (Harterink et al., 2017). The Centrin2-KR fusion protein was generated by cloning the Centrin2 cDNA in-frame with KillerRed, using the BamHI and XhoI cloning sites of the pKillerRed-N vector (Evrogen) (de Anda et al., 2010). P. Gordon-Weeks kindly provided Drebrin-YFP (Addgene plasmid # 40359); (Geraldo et al., 2008) and Drebrin-S142D-YFP (Addgene plasmid S142D Drebrin-YFP # 58336; (Worth et al., 2013). M. Davidson provided Lifeact-mEos3.2 (Addgene plasmid mEos3.2-Lifeact-7 # 54696), mEos3.2 (Addgene plasmid mEos3.2-C1 #54550), Actin-mEos4b (Addgene plasmid mEos4b-Actin-C-18 #57500) and Lifeact-RFP (Addgene plasmid mTagRFP-T-Lifeact-7 #54586) plasmids. F. Bradke (DZNE, Bonn) kindly provided the Lifeact-GFP plasmid. M. Kneussel (ZMNH, UKE) kindly provided the EB3-GFP and X. Chai (from the laboratory of M. Frotscher, ZMNH, UKE) kindly gave us the Cofilin-S3E-GFP construct (Chai et al., 2016). Thomas Oertner (ZMNH, UKE) kindly provided the tDimer (pAAV-CAG-tDimer) plasmid. mMaroon1 (pcDNA3.1-mMaroon1) plasmid was provided by Michael Lin (Addgene plasmid # 83840; http://n2t.net/addgene: 83840; RRID: Addgene_83840). PaGFP-UtrCH was a gift from William Bement (Addgene plasmid # 26738; http://n2t.net/addgene: 26738; RRID: Addgene_26738)

**Animal experiments.**

Animal (rat and mouse) experiments were performed according to the German and European Animal Welfare Act and with the approval of local authorities of the city-state Hamburg (Behörde für Gesundheit und Verbraucherschutz, Fachbereich Veterinärwesen) and the animal care committee of the University Medical Center Hamburg-Eppendorf.

**Preparation of hippocampi and cortices from rat and mouse embryos.**

Pregnant rats and mice were anaesthetized with $CO_2/O_2$ and then euthanized before taking the embryos out from their uteri. Embryos were then decapitated, and heads were collected in petri dishes kept on ice. After opening the skulls, brains were collected in petri dishes with HBSS on ice. Hemispheres were separated, meninges were carefully stripped away, and hippocampi/ cortices were dissected.

**Hippocampal neuronal cultures and transfections.**

Isolated hippocampi (from E18 embryos) were triturated in 1xHBSS (Invitrogen) after digestion by papain and DNase for 10 min at 37°C (Worthington). Transfections were performed using the Amaxa nucleofector system following the manufacturer's manual. 5 x $10^6$ cells and 3 µg of DNA mix were used for each transfection. Lifeact-GFP, Lifeact-RFP, Centrin2-RFP, Lifeact-mEos3.2, mEos3.2, Actin-mEos4b, Drebrin-YFP, Drebrin S142D-YFP, CofilinS3E-GFP, Centrin2-KR, EB3-mCherry, EB3-GFP, tDimer, mMaroon1 and PaGFP-UtrCH plasmids were used for hippocampal neuronal transfections. The final concentration for each plasmid was 1µg, except for Lifeact-GFP, Lifeact-RFP and tDimer. We used 0.5µg of Lifeact-GFP, Lifeact-RFP and 0.3µg of tDimer plasmids. In some cases, empty pcDNA 3.1 was used to make up to 3µg of DNA per each transfection mix as per the manufacturer recommendation. After electroporation, neurons were plated on poly-L-lysine coated glass coverslips (for immunostaining), on glass-bottomed dishes (ibidi, for live imaging) or tissue culture chambers (Sarstedt, for live imaging) in Neurobasal/B27 medium (Invitrogen) and were maintained in culture for 24 hours at 37°C with 5% $CO_2$ before use.

***In utero* electroporation.**

Pregnant C57BL/6 mice with E15 embryos were first administered with pre-operative analgesic, Buprenorphine (0.1 mg / kg), by subcutaneous injection. After 30 min, mice were anaesthetized with isoflurane (4% for induction, 2-3% for maintenance) in oxygen (0.5-0.8 L/min for induction and maintenance). Later, uterine horns were exposed and plasmids mixed with Fast Green (Sigma) were microinjected into the lateral ventricles of embryos. Five current pulses (50 ms pulse / 950 ms interval; 35-36 V) were delivered across the heads of embryos. After surgery, mice were kept in a warm environment and were provided with moist food containing post-operative analgesic, Meloxicam (0.2-1 mg/kg), until they were euthanized for collection of the brains from the embryos. The brains were either used for cortical cultures or cortical slices.

For cortical cultures, we introduced PCM-1 shRNA (with or without PCM-1-GFP) or control shRNA plasmids together with Lifeact-GFP, Lifeact-RFP or Venus expressing plasmids into brain cortices at embryonic day 15 (E15), and isolated cortical neurons at E17. The concentration of shRNA (control or PCM-1 -shRNA), PCM-1-GFP plasmids injected was 3-fold higher than that of the Lifeact-GFP, Lifeact-RFP or Venus plasmids. We used 1.5 µg/ul for shRNA (control or PCM-1 -shRNA),

PCM-1-GFP and 0.5 µg/ul for Lifeact-GFP, Lifeact-RFP and Venus plasmids. Neurons were cultured *in vitro* for an additional 24 hrs and were prepared for time-lapse experiments or pharmacological treatment.

For cortical slices, we introduced Centrin2-RFP together with Lifeact-GFP or PCM-1 shRNA or control shRNA plasmids together with Venus or DeAct-SPvB together with mCherry into brain cortices at embryonic day 15 (E15) and brains were collected at E18. The concentration of Centrin2-RFP, shRNA (control or PCM-1 - shRNA), DeAct-SpvB plasmid injected was 2-3-fold higher than that of the Lifeact-GFP, Venus or mCherry plasmids. We used 1.5 µg/ul of shRNA (control or PCM-1 - shRNA), 1.0 µg/ul for Centrin2-RFP, DeAct-SpvB and 0.5 µg/ul for Lifeact-GFP, Venus and mCherry.

**Cortical cultures.**

Neurons were transfected by *in utero* electroporation at E15 and transfected cortices were dissected two days later (as explained above). Isolated cortices were triturated in 1xHBSS (Invitrogen) containing papain and DNase at 37°C (Worthington; (de Anda et al., 2010). Neurons were plated on poly-L-lysine coated glass-bottomed dishes (ibidi) in Neurobasal/B27 medium (Invitrogen) and were maintained in culture for 24 hours before time-lapse imaging.

**Cortical slices.**

Embryonic brain cortical regions were transfected via *in utero* electroporation at E15. Lifeact-GFP and Centrin2-RFP were used to inspect F-actin distribution, PCM-1 shRNA or DeAct were used to perturb the F-actin organization. The brains collected from E18 embryos were then post fixed in 4% paraformaldehyde (PFA) overnight at 4°C and later moved to 30% sucrose (in PBS) until they were completely sunk. Brains were then embedded in Tissue Tek OCT compound and stored at - 80°C until they were sectioned to 50 µm slices using a cryostat.

***STED microscopy.***

All STED images were acquired on a Leica TCS SP8 gated STED microscope equipped with a pulsed 775 nm depletion laser (80 MHz) and a pulsed white light laser (WLL) for excitation. The microscope was covered by an incubation chamber Black i8 2000 LS (PeCon GmbH, Erbach, Germany) fitted to the Leica microscope stand DMI 6000AFC. The chamber can be temperature controlled with the PeCon Temp Controller 2000-2 and Heating Unit 2000. For acquiring images, Leica Objective HC APO CS2 100x/1.40 Oil was used.

**STED imaging in fixed cells.**

Neurons stained for actin with Atto 647N-Phalloidin (1:40; Stock 10 nM, Sigma-Aldrich, #65906) and pericentrin with anti-rabbit Atto 594 (1:200; Sigma-Aldrich) or anti-rabbit Abberior Star 580 (1:200; Abberior GmbH Gottingen, Germany) were embedded in Mowiol and excited via the WLL at 650 nm and 561 nm, respectively. Emission was acquired between 660-730 nm for Atto 647N and 580-620 nm for Atto 594. The detector time gates for both channels were set from 0.5-1 ns to 6 ns. Both dyes were depleted with 775 nm. Respective confocal channels use the same settings as STED channels, except for a reduction of excitation power. Detection time gates were set from 300 ps to 6 ns for both channels. The Format for all images was set to 1024x1024. Optical zoom of 5 resulted in a voxel size of 23 nm for x-y and 100-160 nm for z. Images where then taken with 600 lines per second and line averaging of 8.

**STED imaging in live cells.**

For live STED-microscopy of DIV 1 hippocampal primary neurons, 250 nM of SiR-actin (tebu-bio GmbH, Offenbach, Germany via Spirochrome AG, Switzerland, (Lukinavicius et al., 2014) were directly added to the conditioned medium and cells were imaged after 5 hours.

Time-lapses (1.7 sec interval) were imaged in pre-carbonated extracellular solution at 37°C. SiR-actin was excited using a 633 nm light from the WLL and fluorescence signals were detected via internal Leica Hybrid Detectors in the range of 650-750 nm. The detection time gate was set to 0.3-6.0 ns. Depletion was done at 775 nm at low depletion power of 30%. Application of a zoom of 5 and a format of 1024x256 resulted in a pixel size in x-y of 23 nm. Scan speed was set to 600 Hz by applying 4 times line averaging.

To improve image quality raw data of STED images were deconvolved using the STED package Huygens Professional (SVI, v 15.10) as follows: To calculate the theoretical point spread function (PSF) the optical microscopic parameters provided with the Leica lif-file were used. Within the *Deconvolution wizard*, time-lapse images were subjected to the automatic background calculation of Huygens software. For deconvolution, the *Signal to noise ratio* was set to 15. The *Optimized iteration mode of the CMLE* was applied, until it reached a *Quality threshold* of 0.05.

***SMLM, Photoconversion and FRAP.***

Photoconversion were performed on a Visitron Systems VisiScope TIRF/FRAP imaging system based on a Nikon Ti-E equipped with a Nikon CFI Apo TIRF 100x,

1.49 NA oil objective, a back focal TIRF scanner for suppression of interference fringes (ILas-2, Roper Scientific France/ PICT-IBiSA, Institut Curie) and controlled via VisiView software. 405, 561, 488 and 647 nm laser lanes were used for illumination and activation of respective fluorophores. Fluorescence was detected through an ET 405/488/561/640 Laser Quad Band filter with The ORCA-Flash 4.0 LT sCMOS camera. Cells were maintained at 37°C in a temperature, CO2 and humidity controlled environmental chamber for all live-imaging experiments.

**SMLM.**

For SMLM imaging of F-actin primary neurons were fixed at DIV1 and immonostained against pericentrin. Phalloidin conjugated to Alexa Fluor 647 (Life Technologies) was added to PBS in a 1:50 dilution and incubated at room temperature (RT) for 3 hours. After extensive washing, samples were post fixed for 10 min with 4% PFS + 4% Sucrose in PBS, washed again with NT buffer (50 mM Tris-HCl pH 8, 10 mM NaCl). Samples were sealed using cavity coverslips filled with Buffer TN containing 10% glucose, 10 mM freshly prepared mercaptoethylamine (pH adjusted to 8 with KOH; Sigma), 40 µg/mL catalase (Sigma), and 0.5 mg/mL glucose oxidase (Sigma).

For SMLM imaging of Alexa Fluor 647, the sample was continuously illuminated with 640 nm light. In addition, the sample was illuminated with 405 nm light at increasing intensity to keep the number of fluorophores in the fluorescent state constant. Between 6000 and 10000 frames were recorded per acquisition with an exposure time of 50 ms. Reconstruction of SMLM images was done as described previously using custom written software (Detection of Molecules; (Yau et al., 2014), (Mikhaylova et al., 2015)).

**Photoconversion experiments.**

Photoconversion of primary rat hippocampal neurons expressing mEos3.2 or mEos4b-tagged constructs was performed using the ILas2 system. Photoconversion was performed using the 405 nm laser with 1% laser power. For the cells in which soma was photoconverted, the area of irradiation was in the range of 520 pixels (2.197 $\mu m^2$) to 1700 pixels (7.2 $\mu m^2$). The cells in which growth cones were photoconverted, the irradiated area was in the range of 2400 pixels (10.1 $\mu m^2$) to 2680 pixels (11.3 $\mu m^2$). The time of 405-laser illumination was always set to 1 ms per pixel (1 pixel = 65 nm). For recording green and red fluorescence, cells were sequentially illuminated with 488 nm laser (2% power) and 561 nm laser (5% power) for 400 ms (300 nm TIRF penetration depth). Three frames each of green and red fluorescence were recorded before and 150 frames (800 ms /frame i.e., 120 sec) each were recorded after the photoconversion. The green and red fluorescence was detected through the 540 nm and 620 nm emission filters respectively.

**Photoactivation experiments**.

Photoactivation of PaGFP signal was performed either in the soma or growth cone of primary rat hippocampal neurons using the ILas2 system. To identify transfected neurons prior to photoactivation, the cultures used for this experiment were always co-transfected with tDimer or mMaroon1 together with PaGFP-UtrCH. To illustrate the neuronal morphology, an image with tDimer or mMaroon1 signal was captured before the photoactivation with 561 nm (3% power with 200 ms exposure time) or 640 nm (10% laser power with 500 ms exposure time) lasers respectively. Photoactivation of PaGFP-UtrCH was achieved using the 405 nm laser with 50% laser power. The photoactivated region was 1240 pixels (5.239 $\mu m^2$); the 405 nm laser illumination time was set to 2 ms per pixel (1 pixel = 65 nm). Cells were illuminated with 488 nm laser (5% power) for 800 ms to record one frame (600nm TIRF penetration depth). Three frames were imaged before and 150 frames (800 ms/frame i.e., 120 sec) were recorded after the photoactivation.

**FRAP experiments**.

Photobleaching of GFP signal was performed either in the soma or growth cone of Lifeact-GFP expressing primary rat hippocampal neurons using the ILas2 system. Photobleaching was achieved using the 405 nm laser with 100% laser power. The photobleached region was 720 pixels (3.04 $\mu m^2$); the 405 nm laser illumination time was set to 1 ms per pixel (1 pixel = 65 nm). Cells were illuminated with 488 nm laser (8% power) for 75 ms to record one frame (70-75 nm TIRF penetration depth). Three frames were imaged before and 188 frames (75 ms/frame i.e., 14.1 sec) were recorded after photobleaching.

**Pharmacological treatments**

Pharmacological compounds were directly added to DIV1 hippocampal neurons in culture. The treated cells were either used for PFA (4%) fixation for immunostaining or time-lapse live imaging. Compounds used and their final concentrations: cytochalasin D (cytoD; actin depolymerizing agent) - 1 μM, jasplakinolide (jasp; actin stabilizing agent) – 300 nM, nocodazole (Noc; microtubule-destabilizing agent) – 7 μM, Brefeldin A – 10μg/ml, SMIFH2 (Formin inhibitor) - 25 μM and CK666 (Arp2/3 inhibitor) - 50 μM. All the compounds listed here were purchased from Sigma.

**Photoconversion/ Photoactivation analysis.**

The 405 nm laser irradiated area for all the cells used for the analysis was between 5.239 and 7.182 $\mu m^2$. 561 nm channel time-lapses of photo-converted cells were preprocessed with ImageJ (Schindelin et al., 2012; Schneider et al., 2012) by subtracting the first empty frame from the remaining frames. For analysis, the area of photoconversion at either soma or growth cones was selected as a ROI. Further ROI's were selected in the corresponding compartments (Growth cone, soma or neurite tip). Average gray-values over time were measured via the Timeseries Analyzer plugin. Initial gray-values in the photoconverted area were normalized to 1, gray-values in the "receiving" compartments were normalized to fractions of initial gray-value at the photoconverted area. The ratio of the average growth cone intensity to the average soma intensity was plotted over time for different experimental groups. For the sake of clarity every four seconds were considered. Furthermore, the intensity ratio of growth cone to soma after two minutes was compared to the intensity ratio of growth cone to soma before photoconversion (green channel). The association of these parameters was determined via Pearson correlation. For the decay of signal in the photoconverted area in the soma t ½ was determined via Graph-Pads fitted one-phase exponential decay equation. The same analysis was performed for photoactivated PaGFP-Utrch expressing cells.

**FRAP analysis**.

Fluorescence recovery after photobleaching in the somatic and growth cone areas of the primary hippocampal neurons expressing Lifeact-GFP was measured using the FRAP profiler and Intensity vs Time Plot plugins (ImageJ). A simple ratio bleach correction was applied before analysis.

**Actin & PCM-1 nearest neighbor analysis.**

Actin- and PCM-1 puncta were counted manually within the three-dimensional soma space and marked in their estimated center with ImageJ's selection tools. Center coordinates were then exported via the ROI manager for further processing with R. For each coordinate a distance matrix was calculated and filtered to find the nearest neighbor. Shortest distances of puncta to the closest object - within and between the actin and PCM-1 datasets - were collected and plotted as a relative frequency distribution. Additionally, the average minimum distance was reported.

**Immunocytochemistry.**

Rat hippocampal neurons grown on coverslips were fixed with 4% paraformaldehyde (PFA) at 37°C for 10 min and then permeabilized with 0.5% triton-x 100 for 10 min. Nonspecific binding was blocked by incubation with 5% donkey serum in PBS for 60 min at RT, followed by specific primary antibody incubation: rabbit anti-pericentrin (Convance/ dcs diagnostics, PRB-432C), rabbit anti-PCM-1 (kindly gifted by Andreas Merdes) or mouse anti-α-tubulin (Abcam, ab7291) was added for incubation for 120 min at RT. The respective anti-mouse or anti-rabbit Alexa fluor -488 or -568 labeled secondary antibody was added for 60 minutes at RT. Primary and secondary antibodies were diluted in PBS with 2% donkey serum. After primary and secondary antibody incubation three washing steps with PBS were performed. For F-actin labeling, cells on coverslips - after PFA fixation and permeabilization - were incubated with Phalloidin 488 from Cytoskeleton Inc. (for confocal imaging) or Phalloidin 647 (for pharmacology experiments, Invitrogen), followed by three PBS washes. In some experiments, Hoechst dye (1:10000, Invitrogen) was added to stain nuclei. Coverslips were mounted onto slides using prolong gold (Invitrogen) and were stored protected from light.

**Epi-fluorescence imaging.**

Epi-fluorescence imaging was performed on an inverted Nikon microscope (Eclipse, Ti) with a 60x objective (NA 1.4). During time-lapse imaging, cells plated on a glass-bottomed dish (Ibidi) or a culture chamber (Sarstedt or Ibidi) were kept in an acrylic chamber at 37°C in 5% $CO_2$. Light intensity of each channel was normally set at 8, with an exposure time of 300-800 ms. Images were captured with a CoolSNAP HQ2camera (Roper Scientific) using NIS-Elements AR software (version 4.20.01 from Nikon Corporation).

**Intensity measurements.**

For CALI experiments growth cone areas were delineated with ImageJ’s selection tools for the first frame of the acquired time-lapse. Mean intensities from these selections were compared for cells before and after CALI treatment. Intensity values after CALI treatment were normalized to intensity values before CALI treatment. For ctrl shRNA, PCM-1 shRNA and PCM-1 rescue, additionally the mean intensity within the soma was measured and the growth cone to soma ratio was calculated.

***Confocal imaging.***

Images were taken using an Olympus FLUOVIEW FV1000 and Zeiss LSM 700 confocal laser-scanning microscopes with 40X objectives (NA 1.3). Z-series images were acquired with a step size of 300 nm or 400 nm.

**Confocal F-actin puncta analysis.**

For centroid analysis shown in Figure 1 A and B, Supplementary figure 1 A, B and Supplementary figure 15, confocal images of DIV1 or DIV2 rat hippocampal neurons labeled with phalloidin (F-actin) and pericentrin antibody were used. The confocal images, with a z-stack spacing of 400 nm) were first thresholded using Auto Local Threshold v1.5 (Image J) with Phansalkar as the method of choice with the radius set according to the size of the cells and their F-actin puncta (radius 2-10, (Neerad et al., 2011)). Despeckle command (ImageJ) was used to reduce noise. The plugin 3D Objects Counter v2.0 (ImageJ, (Bolte and Cordelieres, 2006) was applied on each image for automatic counting of objects as well as for defining the respective centroid coordinates. The counted objects were then manually compared to the original image and false positives were removed. ImageJ's plugin Volumest was used for determination of the cell-volume following the author's instructions (Merzin, Markko. "Applying stereological method in Radiology. Volume measurement." Bachelor Thesis. University of Tartu. 2008). The resulting volume was radially divided from the center of the soma in the xy-plane into 30° segments. F-actin puncta were then assigned to their respective segments based on their position, while the centrosome (stained with pericentrin antibody, shown in red) was always positioned in the first segment in the clock-wise direction. Finally, separate rose plots were generated for all three stages using Microsoft Excel 2010 (Microsoft Corporation) and data was represented as average F-actin puncta/$\mu m^3$. F-actin puncta were further distinguished into cortical and cytosolic. To define F-actin puncta in 10% and 20% distance range of the centrosome, coordinates of cytosolic F-actin puncta were put into perspective of coordinates of the centrosome.

In case of Figure 7 G and H, confocal images of DIV 2 cortical neurons co-transfected with Venus along with control or PCM-1 shRNA plasmids via *in utero* electroporation were used. Confocal images with a z-stack spacing of 300 nm labeled with phalloidin (F-actin) and anti-PCM-1 antibody were used for analysis. Somatic F-actin puncta in control and PCM-1 KD cells were counted manually using ImageJ (NIH). Volumes of the somas were measured using ImageJ (NIH) Volumest plugin as discussed above. Normalized F-actin puncta per $\mu m^3$ somatic volume was plotted for both the groups.

**Analysis of somatic F-actin puncta and F-actin treadmilling in growth cones.**

For analysis of F-actin puncta in the soma and F-actin treadmilling flow in growth cones, 5 min time-lapse videos of Lifeact-GFP or Lifeact-RFP expressing rat hippocampal or mouse cortical neurons were used.

**Analysis of somatic F-actin puncta in living cells.**

For Centroid analysis (shown in Supplementary figure 4 A and B), 5 min time-lapse with 2 sec interval (151 stacks) of the cells that were transfected with Lifeact-GFP and EB3-mCherry were used. Soma area was selected and radially divided into 12 equal segments. In each segment, recognizable puncta were counted through 151 stacks. To characterize puncta blinking preference in the cell body, puncta sum of each segment for all stacks was made and after measuring the area of each segment within cell body, puncta density was calculated and plotted as rose diagrams. Regarding the 'Quadrant' definition, the 3 segments covering the central microtubule organization area (judged based on the co-transfected EB3-mCherry signal) was considered as the first quadrant, namely Q1, followed by Q2, Q3, and Q4 clockwise (each quadrant covers 3 segments accordingly). To examine the blinking frequency of puncta in each quadrant, puncta number of each stack was plotted after converted into percentage by being divided by the total puncta number of all stacks within the same quadrant.

For analysis of stability and total number of F-actin puncta in living cells*,* the soma area of Lifeact transfected neurons from the 5 min time-lapse videos was marked and resliced with an output spacing of 1.0 pixel using ImageJ to create kymographs. Stability duration and total number of F-actin puncta present in the soma area were then analyzed via ImageJ's selection tools, adding selections to the ROI manager. Length of Lifeact labeled structures in the kymographs along the y-axis was used for defining stability duration with each pixel representing 2 seconds (151 frames for 5 min). Based on duration, Lifeact labeled structures were distinguished as unstable puncta: distinct Lifeact spots or vertical lines disappearing in 15 seconds or less; puncta with intermediate stability: vertical Lifeact lines, appearing for 16 to 240 seconds and long-lasting puncta: vertical Lifeact lines that are in the range of 241 to 300 seconds. The total number of puncta per $\mu m^2$ was obtained through the sum of unstable, intermediate and long-lasting F-actin puncta normalized to the area of soma marked for analysis.

**Analysis of F-actin treadmilling in growth cones.**

Kymographs were generated from the growth cones (from the 5 min time-lapse videos) of Lifeact (tagged with GFP or RFP) transfected rat hippocampal or mouse cortical neurons using the ImageJ Kymograph plugin (Code written by J. Rietdorf & A. Seitz, EMBL Heidelberg). From the kymographs (generated by setting line width to 1), the slope of retrograde trajectories of F-actin was measured and average slope was represented in μm/min.

**Laser-induced Chromophore Assisted Light Inactivation (CALI) of Centrin2-KR.**
Rat hippocampal neurons obtained from E18 embryos were co-transfected with Centrin2-KR and either Lifeact-GFP or EB3-GFP and were maintained in culture for 24 hours before experiments. Neurons at stage 2 or early stage 3 were selected. After recording a 5 min time-lapse (2 sec interval) under 100X objective (CFI Apo TIRF, oil, NA 1.49), the neuron selected was irradiated locally at the centrosomal site with FRAP 561 nm laser at 7% laser power for 1.5 s in the region of interest (~2 $\mu m^2$) to inactivate Centrin2-KR and thus to disrupt the centrosome. After 2-3 h of recovery (at 37°C with 5% $CO_2$) the same cells were recorded for another 5 min. Control experiments were performed with the same settings, but cells were irradiated at a somatic site away from the centrosome.

**EB3 comets quantifications.**
For CALI cells, the soma area of EB3-GFP expressing cells was selected. Comets were tracked manually throughout the first two minutes using ImageJ's ROI manager and selection tools. Number of EB3 comets was normalized by area and was reported as average number of comets per $\mu m^2$ * minutes.

***Neurite length measurement and Neurite terminals analysis.***
Cortical neurons co-transfected with Venus and Control or PCM-1-shRNA via in utero electroporation or cells from the Control group treated with cytochalasin D were used for neurite length and neurite terminals analysis. For neurons in situ brains were transfected either with PCM-1-shRNA and venus or DeAct-SpvB and mCherry.

**Neurite length analysis.**
In each cell total length of all neurites, length of the longest neurite and length of other neurites were measured manually from all the three conditions using ImageJ (NIH). In case of in situ analysis only multipolar cells with three or more primary neurites were considered and traced via ImageJ's simple neurite tracer (Longair et al., 2011) to account for the 3D distribution.

Number of primary neurites with length greater than 60 μm per cell was counted manually from all the three conditions.

**Neurite terminals analysis**.

Total number of neurite terminals per cell was counted manually from all the three conditions.

**Image processing.**

Linear adjustment of brightness and contrast was performed on images using Photoshop CS or ImageJ.

**Statistical analysis.**

Statistical analysis was performed using the GraphPad Prism 6 software. Data shown in the graphs were collected from at least three independent experiments. The Student's t-test (two-tailed) was used to compare means of two groups, whereas analysis of variance (ANOVA) was used when comparing more than two groups. Asterisks *, **, *** and **** represent $p < 0.05$, 0.01, 0.001 and 0.0001 respectively. Error bars in the graphs always represent standard error of mean. Pearson correlation analysis was performed for showing a linear relationship between two sets of data.

**Author contributions.**

F.C. de A. conceived the idea, supervised the project, and wrote the manuscript. F.C. de A. and D.P.M. designed the study. B.S., B.Z. and D.P.M. conducted all the cell culture. O.K., M.K., M.M. and D.P.M. performed STED imaging. M.M. carried out the SMLM imaging. D.E. helped with the initial SMLM imaging. D.P.M. and S.K. performed photoconversion, photoactivation and FRAP experiments; R.S. and F.C. de A. analyzed the data. D.P.M. and M.R. performed all *in utero* electroporation surgeries, D.P.M., R.S., and I.S. analyzed the data. B.Z. and S.K. performed CALI experiments, B.Z. and R.S. analyzed the data. R.S. and D.P.M. performed all the confocal imaging. T.K., I.S., and D.P.M. performed F-actin puncta analysis. B.Z. and D.P.M. performed the epifluorescence time-lapse imaging. D.P.M., R.S. B.Z., and S.W. analyzed the data. F.C. de A. and C.G.D. performed pharmacological experiments. F.C. de A. supervised D.P.M., B.Z., R.S., T.K., I.S., B.S., M.R., and M.M. supervised S.K. All authors helped writing the manuscript.

**Fig. S1. Confocal microscopy reveals cytosolic F-actin puncta around the centrosome in developing neurons. (A)** Early stage 3-hippocampal neuron labelled with phalloidin and Pericentrin antibody. Confocal z-stacks from the inset show F-actin puncta around the centrosome. (**B**) Rose plots depict distribution of F-actin puncta (cortical and cytosolic or only cytosolic) in the cell body with respect to the position of the centrosome. 10% and 20% distance range from the total area is indicated in different shades of blue; n=10 cells, obtained from at least three different cultures. (**C**) Bipolar cell located in the IZ/CP of the developing cortex expresses Lifeact-GFP and Centrin2-RFP and shows F-actin puncta surrounding the centrosome. (**D**) Confocal (CLSM) and STED microscopic images of an early stage 3 hippocampal neuron. Inset: STED Z-stack images with 100 nm Z-spacing showing F-actin puncta localizing near the centrosome. Insets from arrowheads in Z-stack images show F-actin puncta with F-actin fibers attached. Scale bar: 10 μm (A, C and D).

**Fig. S2. Super-resolution microscopy reveals cytosolic F-actin puncta with filaments in developing neurons.** (**A**) Epi-fluorescence (EFM) and SMLM (STORM) images of stage 2 neuron. Inset 1: F-actin puncta near the centrosome forming a pocket-like structure. Arrowheads from insets 2 and 3 show individual F-actin puncta with F-actin fibers attached. Scale bar: 10 μm.

**Fig. S3. Higher expression of Lifeact stabilizes F-actin.** (**A**) Cells expressing high levels of Lifeact show less dynamic growth cones (kymographs) compared with cells expressing Lifeact levels similar to the levels detected with phalloidin staining (B and C, respectively). (**B**) Cells expressing Lifeact and labeling F-actin at comparable levels of Phalloidin staining (**C**) have dynamic growth cones (Kymograph in bottom panel of **B**). (**D**) Enlarged images of the soma from cells shown in A-C. Scale bar: μm (A, B and C).

**Fig. S4. Dynamic F-actin puncta organized around the MTOC and show different blinking frequencies.** (**A**) Maximum projection of a time-lapse of stage 1 neuron expressing EB3-mCherry and Lifeact-GFP. Higher density of F-actin puncta co-localize in quadrant 1 (Q1) with more more EB3 comets. (**B**) Left panel: distribution of F-actin puncta in the cell body from stage 2 cells and stage 3 cells. Right panel: percentage of F-actin puncta per quadrant during time-lapse analysis from stage 2 cells and stage 3 cells. The MTOC is always positioned at quadrant 1(Q1); $p<0.0001$, all by one-way ANOVA, *post hoc Dunnett test*; ***$p<0.001$, ****$p<0.0001$. Mean ± SEM; n=10 cells for each stage, from at least three different cultures. (**C**) Neuron (DIV1) transfected with Lifeact-GFP showing somatic F-actin puncta. (**D**) Montage and kymograph generated from the time-lapse of cell shown in C (region marked with red square), show dynamic F-actin punctum. Red arrowheads in the kymograph show comets generated from the blinking F-actin punctum. (**E**) Duration of F-actin puncta in the cell body of developing neurons. Unstable puncta (% in the somatic area) = 90.13 ± 1.33; Intermediate stability puncta = 9.64 ± 1.31; Long lasting puncta = 0.22 ± 0.03. Mean ± SEM; n=20 cells, from at least three different cultures. Scale bar: 5 μm (A, C)**.**

**Fig. S5. Super-resolution microscopy reveals that higher concentration of SiR-actin (500nM) induced the formation of somatic F-actin fibers**. (**A**) With a higher concentration of SiR-actin, the somatic F-actin puncta have longer F-actin fibers attached to them with less activity (lower panel arrowheads), compared with the lower SiR-actin concentration used (Figure 1E). Scale bar: 2 μm and 0.5 μm (insets).

**Fig. S6. Somatic F-actin turnover is similar to the turnover in growth cones.** (**A**) Lifeact-GFP expressing stage 2 hippocampal neuron photobleached in the soma (region marked by red circle) using 405 nm laser. The kymograph obtained from the photobleached region illustrates fluorescence recovery of the F-actin punctum after photobleaching. (**B**) Time course of the normalized fluorescence intensity in FRAPed region in the somatic and growth cone regions of Lifeact-GFP expressing neurons. Half-time (t ½) values were shown in the inset. t ½ (in sec) for somatic puncta = 1.584 ± 0.2151, growth cones = 2.030 ± 0.2422. n.s = not significant, p=0.2224 by unpaired Student's t test. Mean ± SEM; n=9 cells for somatic puncta and n=15 cells for growth cones. Cells were obtained from at least two different cultures. Scale bar: 10 μm (A)

**Fig. S7. Comparison of the somatic photoconvesion of Lifeact, Actin and Eos alone.** (**A**) Lifeact-mEos3.2, (**B**) Actin-mEos4b, and (**C**) mEos3.2 expressing cells photoconverted in the soma using 405 nm laser (red circle). Cell before and after photoconversion show probe-specific behavior. Arrowheads point the reach of the photoconverted signal over time. Scale bar: 10 μm.

**Fig. S8. F-actin disruption affects release of somatic F-actin from the puncta to the periphery in developing neurons.** (**A**) Time-lapse analysis revealed that F-actin cluster formation following Cytochalasin D treatment originated from pre-existing intermittent F-actin puncta (7 cells from three different cultures). (**B**) Cytochalasin D treatment produced F-actin clusters around the centrosome (83.44%; n=157 from at least three different cultures). (**C**) Brefeldin A treatment does not affect F-actin clusters after Cytochalasin D treatment (95.24%; n=21 cells from at least three different cultures). (**D**) Lifeact-mEos3.2 expressing cell treated with Cytochalasin D (1 μM) and photoconverted in the soma with 405 nm laser (red circle). Cell before (green) and after (red) photoconversion. Red arrowheads point the reach of the photo converted signal over time. (**E**) Cytochalasin D treatment (1 μM) precludes translocation of somatic photoconverted Lifeact-mEos3.2 signal to the cell periphery (ratio neurite tip/ soma). Experiments shown in Figure 6 C, D, Supplementary figure 8 D, E and Supplementary figure 10 C, D were done at the same time, therefore the same Control data is used for comparison. Mean ± SEM, n = 12 untreated cells, n = 8 cells for cytochalasin D from at least 3 different cultures. Scale bar: 5 μm (A, B) and 10 μm (C).

**Fig. S9. Photoconverted F-actin (Lifeact-mEos3.2) in growth cones does not translocate towards the cell body.** (**A**) Lifeact-Eos3.2 (**B**) Actin-mEos4b, and (**C**) mEos3.2 expressing cells photoconverted in the growth cones using 405 nm laser (red circle). Cells before and after photoconversion. Red arrowheads point the reach of the photoconverted signal over time. Scale bar: 10 μm (A, B and C).

**Fig. S10. Jasplakinolide treatment blocks F-actin at the soma.** (**A**) Jasplakinolide promotes the formation of a F-actin pocket-like structure in the soma (91.01% of 189 cells from at least three different cultures). (**B**) Neuron treated with Jasplakinolide shows the formation of a F-actin structure at the region of higher EB3 density. Kymograph shows that this F-actin structure is not dynamic. (**C**) Jasplakinolide (300 nM) treated stage 2 Lifeact-mEos3.2 expressing cell photoconverted in the soma with 405 nm laser (red circle). Cell before (green) and after (red) photoconversion. Red arrowheads point the reach of the photo-converted signal over time. (**D**) Jasplakinolide treatment (300 nM) precludes translocation of somatic photoconverted Lifeact-mEos3.2 signal to the cell periphery (ratio neurite tip/ soma). Experiments shown in Figure 6 C, D, Supplementary figure 8 D, E and Supplementary figure 10 C, D were done at the same time, therefore the same Control data is used for comparison. Mean ± SEM, n=12 untreated cells and n=6 cells for Jasplakinolide from at least 3 different cultures. Scale bar: 10 μm.

**Fig. S11. Reducing the dynamics of F-actin, by overexpressing Drebrin or Cofilin phospho-mimetic mutant, unveils F-actin release from the puncta to the periphery.** (**A**) Representative growth cone from stage 2 neurons and respective kymographs obtained from 5 min time-lapses show F-actin treadmilling from control (Lifeact-GFP; 5.130 ± 0.1070 μm per min, n=15 from at least three different cultures), Drebrin-S142D (Drebrin-S142D-YFP+Lifeact-RFP; 2.080 ± 0.06728 μm per min, n=10 from at least three different cultures; (Zhao et al., 2017)) and Cofilin-S3E (Cofilin-S3E-GFP+Lifeact-RFP; 1.772 ± 0.07147 μm per min, n=10 from at least three different cultures) groups. (**B**) Representative stage 2 cell transfected with Lifeact-RFP and Cofilin-S3E-GFP. Middle panel: Time-lapse montage (from Inset in H) and kymograph (right panel) obtained from green channel shows F-actin sliding through the pre-existing F-actin fiber and initiating the formation of growth cone like protrusion (pointed out by red arrowheads). Scale bar: 2 μm (A) and 10 μm (B).

**Fig. S12. CALI in somatic region distant from Centrosome does not affect F-actin intensity and treadmilling in growth cones.** (**A**) Neurons transfected with Centrin2-KillerRed and EB3-GFP subjected to CALI. Left: Max projection of time-lapse (40sec) before treatment; Right: Max projection of time-lapse (40sec) after treatment. (**B**) Number of EB3 trajectories in the soma of example cell over time (before and after treatment). EB3 trajectories per $\mu m^2$ and minute compared by paired Student's t-test. Before CALI: 2.998 ± 0.279; after CALI: 2.171 ± 0.330; Mean ± SEM; n=7 cells from three different cultures. (**C**) Neurons transfected with Centrin2-KR and Lifeact-GFP were subjected to CALI in somatic regions distant from centrosome (control region). Left panel: Centrin2-KR signal before and after CALI; Middle and Right panels show Centrin2-KR labeling the centrosome before and after CALI and irradiated region marked as a red circle. Enlarged images of the growth cones before (marked as 1) and after (marked as 2) CALI, red lines indicate the areas from where respective kymographs of actin treadmilling were obtained. (**D**) Control irradiation does not affect F-actin treadmilling rate and F-actin intensity in growth cones. Mean ± SEM; n=10 cells from at least three different cultures; p = 0.1522 by paired Student's t-test. Scale bar: 5 μm (A) 10 μm (C).

**Fig. S13. PCM-1 intermingles with F-actin puncta.** (**A**) PCM-1-GFP expressing-cells show close association of PCM-1 and F-actin puncta. Kymographs obtained from white-line and arrowheads in Max Projection of the time-lapse show close association of F-actin and PCM-1. (**B**) Jasplaklinolide treatment induces the formation of a somatic F-actin "ring" structure accompanied by PCM-1 particles**.** (**C**) Cytochalasin D treatment leads to the formation of somatic F-actin clusters accompanied by PCM-1 particles. (**D**) Nocodazole treatment in combination with Cytochalasin D disperses somatic F-actin clusters which are accompanied by PCM-1 particles. Cells were stained with PCM-1 antibody, phalloidin (labels F-actin) and DAPI (stains nucleus). Scale bar: 10 μm (A), 5 μm (B-D).

**Fig. S14. PCM-1 downregulation affects neurite elongation.** (**A**) DIV 2 cortical neurons from E17 mouse embryos that were in utero electroporated at E15 with Control or PCM-1-shRNA or cells from the Control group were treated with 1μM Cytochalasin D 24 h after plating for Cytochalasin D condition. Venus was co-electroporated in all the conditions as a transfection marker. Anti-PCM-1 antibody staining shown in the PCM-1 shRNA insert confirms efficient knockdown of PCM-1 in the Venus transfected cell (pointed with a red arrow). (**B**-**D**) Cytochalasin D and PCM-1 down-regulation boost similarly neurite outgrowth. (**B**) Total length of neurites per cell (in μm). Length of all neurites in Control condition = 269.5 ± 12.18, PCM-1-shRNA condition = 329.5 ± 12.55, cytochalasin D condition = 303.3 ± 15.38. Length of longest neurite in Control condition = 117.8 ± 8.054, PCM-1-shRNA condition = 184.0 ± 12.05, cytochalasin D condition = 190.3 ± 13.06. Length of other neurites in Control condition = 149.5 ± 11.58, PCM-1-shRNA condition = 145.4 ± 9.357, cytochalasin D condition = 113.0 ± 7.318. $p<0.0001$ by one-way ANOVA, post hoc Tukey test, ***$p<0.001$, **$p<0.01$, n.s = not significant. (**C**) Number of neurites greater than 60 μm per cell in Control condition = 1.120 ± 0.1093, PCM-1-shRNA condition = 1.520 ± 0.1040, cytochalasin D condition = 1.980 ± 0.1414; $p<0.0001$ by one-way ANOVA, post hoc Dunnett's test, ****$p<0.0001$, *$p<0.05$. (**D**) Quantification showing number of neurite tips per cell in Control condition = 6.30 ± 0.3571, PCM-1-shRNA condition = 8.780 ± 0.52, cytochalasin D condition = 6.10 ± 0.3651; $p<0.0001$ by one-way ANOVA, post hoc Dunnett's test, ***$p<0.001$. Mean ± SEM; n=50 cells for each group from at least three different cultures (for data shown in **B**, **C** and **D)**, Scale bar: 10 μm (A)

**Fig. S15. Somatic F-actin puncta are Formin but not Arp2/3 dependent.** DIV 1 neurons incubated for two hours with (**A**) DMSO or (**B**) 50 μM CK666, (**C**) 25 μM SMIFH2 (**D**) 50 μM CK666 + 25 μM SMIFH2 stained with anti-pericentrin antibody and Phalloidin. (**E**) Cytosolic F-actin puncta distribution from the indicated treatment groups. The proximity of cytosolic F-actin puncta with centrosome is depicted in B (10% or 20% distance range from the total area) (**F**) Total number of cytosolic F-actin puncta per $\mu m^3$ from cells treated with DMSO = 1.000 ± 0.2559, CK666 = 0.7740 ± 0.0693, SMIFH2 = 0.3308 ± 0.0605 and CK666+SMIFH2 = 0.4067 ± 0.0554. $p=0.0036$ by one-way ANOVA, post hoc Dunnet test; *$p<0.05$, **$p<0.01$. Mean ± SEM; n=11 for SMIFH2, n=10 cells each for DMSO, CK666 and SMIFH2+CK666 groups from at least two different cultures (for data shown in **E** and **F**). Scale bar: 5 μm.

**Fig. S16. The effect of PCM-1 down-regulation is reversed by a PCM-1 shRNA resistant plasmid.** (**A**) DIV 1 cortical neuron from E17 mouse embryos *in utero* electroporated at E15 with PCM-1 shRNA together with Lifeact-RFP and Chicken-PCM-1-GFP (PCM-1 shRNA resistant plasmid). Arrowhead points to PCM-1-GFP puncta. Kymographs obtained from white lines marked as 1 (for soma) and 2 (for growth cone). (**B-D**) Quantifications from control (control shRNA + Lifeact-RFP) and PCM-1 rescue (PCM-1 shRNA + Chicken-PCM-1-GFP + Lifeact RFP) conditions: (**B**) Density of somatic F-actin puncta of stage 2 cells from control condition = 1.000 ± 0.06468, PCM-1 rescue condition = 0.8937 ± 0.05249; p = 0.2378 by unpaired Student's t test. Mean ± SEM; n=5 cells each for control and PCM-1 rescue groups. Cells were obtained from at least two different cultures. (**C**) F-actin treadmilling speed (in µm/min) in the growth cones of stage 2 cells from control condition = 4.753 ± 0.1390, PCM-1 rescue condition = 4.905 ± 0.1346, p = 0.4357 by unpaired Student's t test. Mean ± SEM; n=9 cells each for control and PCM-1 rescue groups. Cells were obtained from at least two different cultures. (**D**) F-actin intensity ratio in the growth cones of stage 2 cells from control condition = 1.182 ± 0.05147, PCM-1 rescue condition = 1.113 ± 0.05770. n.s = not significant (p = 0.3746) by unpaired Student's t-test. Mean ± SEM; n=9 cells for control and n=8 cells for PCM-1 rescue from at least two different cultures. Scale bar: 10 µm.

**Movie S1. Time-lapse imaging of somatic F-actin puncta labeled with SiR-actin.** STED imaging was performed on a Leica TCS SP8 gated STED microscope equipped with a pulsed 775 nm depletion laser (80 MHz) and a pulsed white light laser (WLL) for excitation. Duration of time-lapse imaging: 1.5 min. Interval between the frames is 1.7 sec.

**Movie S2. Photoconversion in the soma of stage 2/3 neurons expressing Lifeact-mEos3.2.** Imaging was performed on a Visitron Systems VisiScope TIRF/FRAP imaging system based on a Nikon Ti-E equipped with a Nikon CFI Apo TIRF 100x, 1.49 NA oil objective. Duration of time-lapse imaging: 144 sec; 2.4 sec before and 141.6 sec after photoconversion. Interval between the frames is 0.8 sec.

**Movie S3. Photoconversion in the soma of stage 2 neurons expressing PaGFP-UtrCH and tDimer.** Imaging was performed on a Visitron Systems VisiScope TIRF/FRAP imaging system based on a Nikon Ti-E equipped with a Nikon CFI Apo TIRF 100x, 1.49 NA oil objective. Duration of time-lapse imaging: 120.6 sec; 0.6 sec before and 120 sec after photoactivation. Interval between the frames is 0.2 sec.

**Movie S4. Photoconversion in the soma of stage 2 neurons expressing either Actin-mEos4b or mEos3.2.** Imaging was performed on a Visitron Systems VisiScope TIRF/FRAP imaging system based on a Nikon Ti-E equipped with a Nikon CFI Apo TIRF 100x, 1.49 NA oil objective. Duration of time-lapse imaging: 144 sec; 2.4 sec before and 141.6 sec after photoconversion. Interval between the frames is 0.8 sec.

**Movie S5. Photoconversion in the soma of stage 2 neurons expressing Lifeact-mEos3.2 treated with 1 µM cytochalasin D.** Imaging was performed on a Visitron Systems VisiScope TIRF/FRAP imaging system based on a Nikon Ti-E equipped with a Nikon CFI Apo TIRF 100x, 1.49 NA oil objective. Duration of time-lapse imaging: 144 sec; 2.4 sec before and 141.6 sec after photoconversion. Interval between the frames is 0.8 sec.

**Movie S6. Photoconversion in the growth cones of stage 2 neurons expressing Lifeact-mEos3.2 or Actin-mEos4b or mEos3.2.** Imaging was performed on a Visitron Systems VisiScope TIRF/FRAP imaging system based on a Nikon Ti-E equipped with a Nikon CFI Apo TIRF 100x, 1.49 NA oil objective. Duration of time-lapse imaging: 144 sec; 2.4 sec before and 141.6 sec after photoconversion. Interval between the frames is 0.8 sec.

**Movie S7. Time-lapse imaging of neurons expressing Lifeact-RFP and Drebrin-YFP or Drebrin S142D-YFP.** Epi-fluorescence imaging was performed on an inverted Nikon microscope (Eclipse, Ti) with a 60x objective (NA 1.4). Duration of time-lapse imaging: 5 min. Interval between the frames is 2 sec.

**Movie S8. Time-lapse imaging of neuron expressing Lifeact-RFP and Cofilin S3E-GFP.** Epi-fluorescence imaging was performed on an inverted Nikon microscope (Eclipse, Ti) with a 60x objective (NA 1.4). Duration of time-lapse imaging: 5 min. Interval between the frames is 2 sec.

**Movie S9. CALI of centrosome in early stage 3 neuron expressing Centrin2-KR and EB3-GFP.** Cell was imaged for 5 min (interval between the frames is 2 sec) before laser irradiation (Before CALI). 2-3 hr after irradiation (After CALI) the same cell was imaged 5 min (interval between the frames is 2 sec). Epi-fluorescence imaging was performed on an inverted Nikon microscope (Eclipse, Ti) with a 100x objective (NA 1.49).

**Movie S10. CALI of centrosome in early stage 3 neuron expressing Centrin2-KR and Lifeact-GFP.** Cell was imaged for 5 min (interval between the frames is 2 sec) before laser irradiation (Before CALI). 2-3 hrs after irradiation (After CALI) the same cell was imaged 5 min (interval between the frames is 2 sec). Epi-fluorescence imaging was performed on an inverted Nikon microscope (Eclipse, Ti) with a 100x objective (NA 1.49).

**Movie S11. CALI of somatic region outside of the centrosome in early stage 3 neuron expressing Centrin2-KR and Lifeact-GFP.** Cell was imaged for 5 min (interval between the frames is 2 sec) before laser irradiation (Before CALI). 2-3 hrs after irradiation (After CALI) the same cell was imaged 5 min (interval between the frames is 2 sec). Epi-fluorescence imaging was performed on an inverted Nikon microscope (Eclipse, Ti) with a 100x objective (NA 1.49).

**Movie S12. Photoconversion in the soma of stage 2 neuron expressing Lifeact-mEos3.2 treated with 7 μM nocodazole.** Imaging was performed on a Visitron Systems VisiScope TIRF/FRAP imaging system based on a Nikon Ti-E equipped with a Nikon CFI Apo TIRF 100x, 1.49 NA oil objective. Duration of time-lapse imaging: 144 sec; 2.4 sec before and 141.6 sec after photoconversion. Interval between the frames is 0.8 sec.

**Movie S13. Time-lapse imaging of neuron expressing Lifeact-RFP and PCM-1-GFP.** Epi-fluorescence imaging was performed on an inverted Nikon microscope (Eclipse, Ti) with a 60x objective (NA 1.4). Duration of time-lapse imaging: 5 min. Interval between the frames is 0.8 sec 2 sec.
